# *Arabidopsis* and *Chlamydomonas* phosphoribulokinase crystal structures complete the redox structural proteome of the Calvin-Benson cycle

**DOI:** 10.1101/422709

**Authors:** Libero Gurrieri, Alessandra Del Giudice, Nicola Demitri, Giuseppe Falini, Nicolae Viorel Pavel, Mirko Zaffagnini, Maurizio Polentarutti, Pierre Crozet, Christophe H. Marchand, Julien Henri, Paolo Trost, Stéphane D. Lemaire, Francesca Sparla, Simona Fermani

## Abstract

In land plants and algae, the Calvin-Benson (CB) cycle takes place in the chloroplast, a specialized organelle in which photosynthesis occurs. Thioredoxins (TRXs) are small ubiquitous proteins, known to harmonize the two stages of photosynthesis through a thiol-based mechanism. Among the 11 enzymes of the CB cycle, the TRX target phosphoribulokinase (PRK) has yet to be characterized at the atomic scale. To accomplish this goal, we determined the crystal structures of PRK from two model species: the green alga *Chlamydomonas reinhardtii* (*Cr*PRK) and the land plant *Arabidopsis thaliana* (*At*PRK). PRK is an elongated homodimer characterized by a large central β-sheet of 18 strands, extending between two catalytic sites positioned at its edges. The electrostatic surface potential of the catalytic cavity has both a positive region suitable for binding the phosphate groups of substrates and an exposed negative region to attract positively charged TRX-f. In the catalytic cavity, the regulatory cysteines are 13 Å apart and connected by a flexible region exclusive to photosynthetic eukaryotes—the clamp loop—which is believed to be essential for oxidation-induced structural rearrangements. Structural comparisons with prokaryotic and evolutionarily older PRKs revealed that both *At*PRK and *Cr*PRK have a strongly reduced dimer interface and increased number of random coiled regions, suggesting that a general loss in structural rigidity correlates with gains in TRX sensitivity during the molecular evolution of PRKs in eukaryotes.

**Significance Statement:** In chloroplasts, five enzymes of the Calvin-Benson (CB) cycle are regulated by thioredoxins (TRXs). These enzymes have all been structurally characterized with the notable exception of phosphoribulokinase (PRK). Here, we determined the crystal structure of chloroplast PRK from two model photosynthetic organisms. Regulatory cysteines appear distant from each other and are linked by a long loop that is present only in plant-type PRKs and allows disulfide bond formation and subsequent conformational rearrangements. Structural comparisons with ancient PRKs indicate that the presence of flexible regions close to regulatory cysteines is a unique feature that is shared by TRX-dependent CB cycle enzymes, suggesting that the evolution of the PRK structure has resulted in a global increase in protein flexibility for photosynthetic eukaryotes.

## INTRODUCTION

Photosynthetic CO2 fixation supports life on earth and is a fundamental source of food, fuel, and chemicals for human society. In the vast majority of photosynthetic organisms, carbon fixation occurs via the Calvin-Benson (CB) cycle (Fig. S1), a pathway that has been extensively studied at the physiological, biochemical, and structural levels. The CB cycle consists of 13 distinct reactions catalyzed by 11 enzymes (1) that are differentially regulated to coordinate the two stages of photosynthesis—electron transport and carbon fixation—and to couple them with the continuous changes in environmental light.

Thioredoxins (TRXs) are small thiol oxidoreductase enzymes that are widely distributed among the kingdoms of life. They reduce disulfide bonds in target proteins, controlling the redox state and modulating the activity of target enzymes (2). Compared to animals that usually possess 2–3 different TRXs, the plant thioredoxin system is much more complex. Looking only at the chloroplast, five different classes of TRXs are known (i.e., TRX-f, -m, -x, -y, and -z) with several isoforms composing many of these classes (2). Despite this diversity, TRXs are ancient and strongly evolutionarily conserved, consisting of a central core of a five-stranded β-sheet surrounded by four α-helices and an active site that protrudes from the protein’s surface (3).

In TRX-dependent regulation, a small fraction of photosynthetic-reducing equivalents are transferred from photosystem I to several chloroplast TRXs, which in turn reduce key enzymes of the CB cycle, regulating their activity (1). Among the five classes of chloroplast TRXs, class f displays a high specificity towards CB cycle enzymes. In both plants and algae, the activities of fructose-1,6-bisphosphatase (FBPase) (4, 5), sedoheptulose-1,7-bisphosphatase (SBPase) (5), PRK (6), and the AB isoform of NADP-glyceraldehyde 3-phosphate dehydrogenase (AB-GAPDH) (7) in plants and of phosphoglycerate kinase (PGK) (8) in *Chlamydomonas* are directly linked to light by a thiol-based mechanism driven by TRX (Fig. S1). In addition, PRK forms a regulatory complex with the redox-sensitive CP12 and the homotetrameric NADP-GAPDH (A_4_-GAPDH) (9–11). The formation of this ternary complex occurs in both plants and algae, and causes an almost complete inhibition of PRK activity (12).

In contrast to the canonical WCGPC active site of TRXs, no thioredoxin-recognition signature is found in these enzymes. A structural comparison of SBPase and FBPase highlights how evolution has adopted different strategies for modeling protein structure to achieve the same TRX-dependent redox control (5). Of the three enzymes regulated by TRX whose 3D structures are available, a pair of redox-sensitive cysteines was found to be located in completely different regions. In AB-GAPDH, these cysteines are located in a C-terminal extension (CTE) (7); in SBPase, they are present at the dimer interface (5); and in FBPase, they are solvent exposed and distant from the sugar-binding site (4, 5).

To complete the structural redox proteome of the CB cycle enzymes and identify the common feature behind the acquisition of TRX-mediated redox control, the crystal structures of PRK from two model species—the green alga *Chlamydomonas reinhardtii* and the land plant *Arabidopsis thaliana*—were determined and compared to those of other TRX-controlled enzymes. Our results strongly show that the acquisition of a TRX-targeted motif positively correlates with the acquisition of flexibility in the target proteins, consistent with the observation that the entropic contribution obtained from the reduction of the oxidized target enzyme is the main driving force of the redox control mediated by TRXs (13).

## RESULTS AND DISCUSSION

### Three-dimensional structures of *Cr*PRK and *At*PRK in solution and in crystal form

To gain insight into the molecular mechanism of regeneration of the five-carbon sugar ribulose-1,5-bisphosphate (Fig. S1) and of its redox control by dithiol/disulfide exchange, the crystal structures of *Cr*PRK and *At*PRK were determined at 2.6 Å and 2.5 Å, respectively, and compared with those obtained in solution by size-exclusion chromatography-small angle X-ray scattering (SEC-SAXS). Both enzymes are dimers of two identical monomers connected by a 2-fold noncrystallographic axis (Fig. 1). They are structurally similar, and their superimposition results in a root main square deviation of 0.61 Å (on 330 aligned residues) for *Cr*PRK and 1.03 Å (on 328 aligned residues) for *At*PRK.

**Fig. 1.**
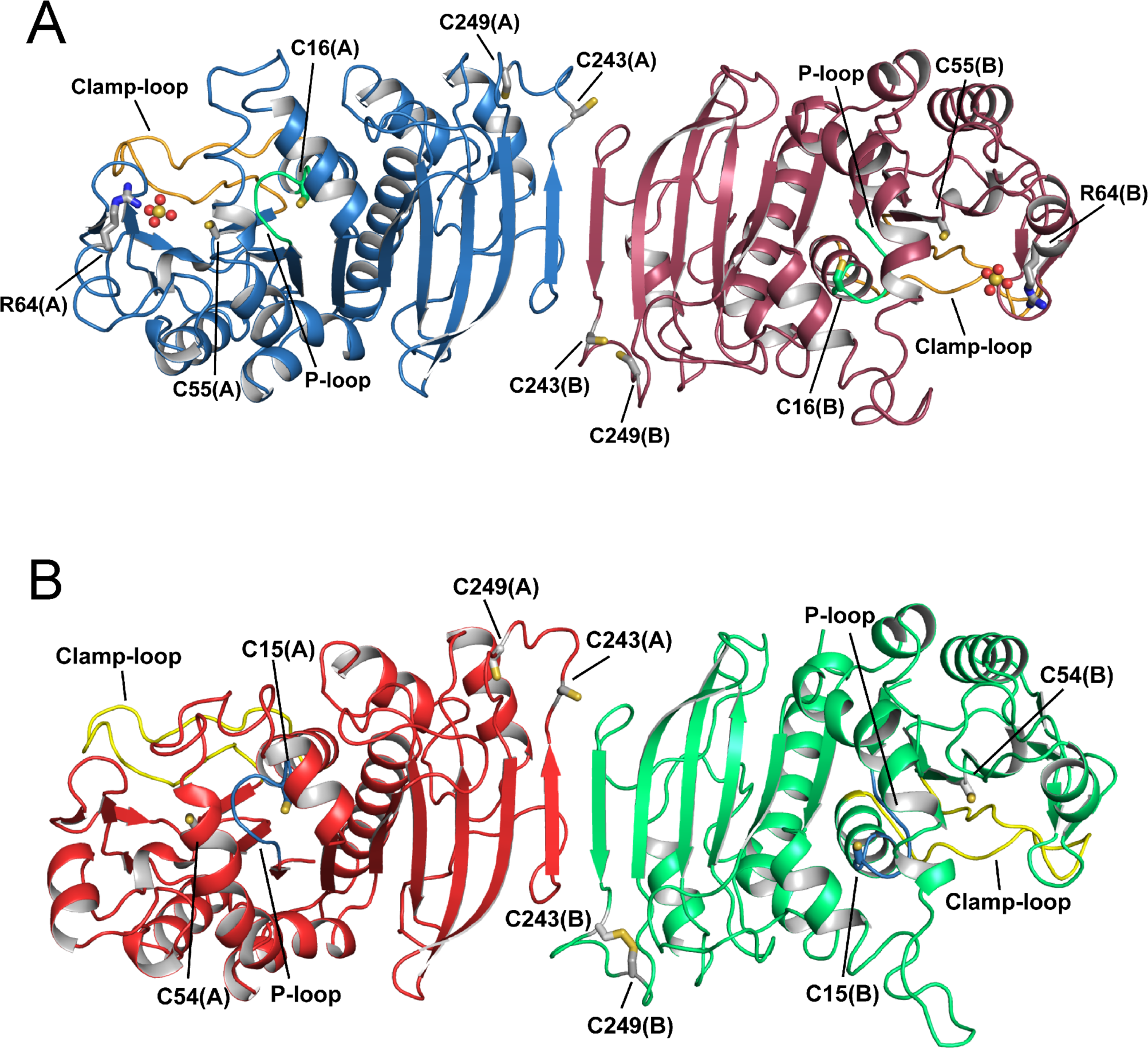
The crystal structure of photosynthetic PRK. Representation of the overall crystal structure of reduced phosphoribulokinase (PRK) from (*A*) *Chlamydomonas reinhardtii* and (*B*) *Arabidopsis thaliana*. The P-loop is highlighted in green and light-blue, and the clamp loop in orange and yellow, respectively. Each monomer contains two pairs of cysteines, one in the active site (Cys16 and Cys55 for *Cr*PRK and Cys15 and Cys54 for *At*PRK) and one in the dimer interface close to the C-terminal end of the protein chain (Cys243 and Cys249). Cysteine residues are indicated and represented as sticks. In *At*PRK, a pair of C-terminal cysteines forms a disulfide bond. *Cr*PRK binds two sulfate ions (one for each monomer), represented as spheres, from the crystallization solution. Arg64, represented as sticks, is one of the residues stabilizing the anions.

Each monomer contains a central nine-stranded mixed β-sheet (β1−β9) surrounded by eight α-helices (α1, α3–α9), four additional β-strands (β1’–β4’), and one additional α-helix (α2; Figs. 2 and S2).

**Fig. 2.**
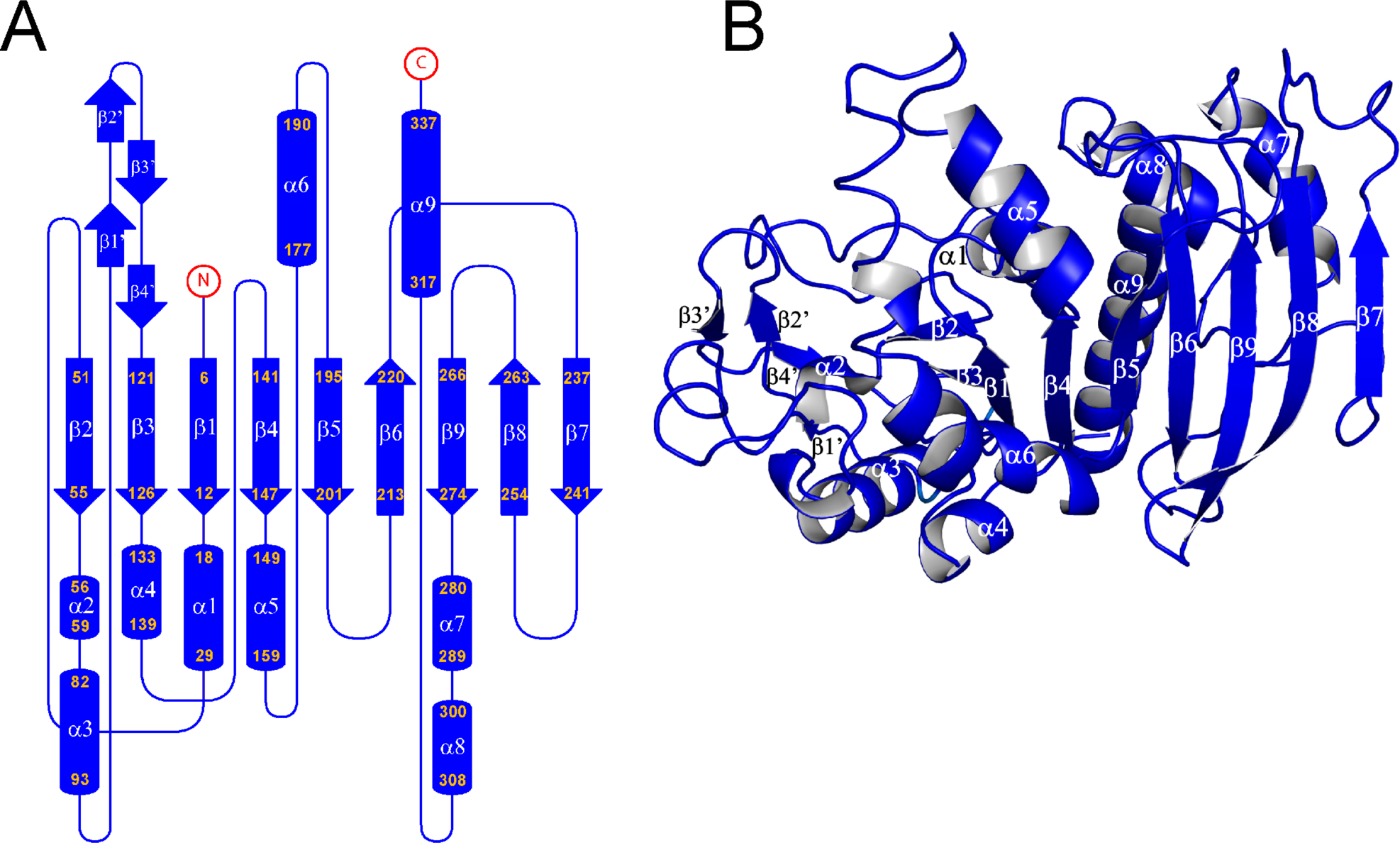
The crystal structure of the *Cr*PRK monomer. (*A*) Topology diagram: The monomer is composed of a mixed β-sheet of nine strands, nine α-helices, and four additional small β-strands indicated by β’. (*B*) Representation of the structure of the monomer: The central β-sheet is sandwiched between helices α3, α4, and α6 and helices α1, α7, α8, and α9. Strand β7 is located in the dimer interface, whereas the four additional β-strands (β1’ to β4’) form the edge of the dimer.

The *At*PRK crystal structure confirms the elongated shape of the oxidized protein previously observed by SAXS analysis (10) but appears less bent and twisted. Structural information regarding *Cr*PRK in solution provides a single recognizable species with an R_g_ value of 35 ± 1 Å and an estimated molecular weight around 70 kDa (Fig. S3 and Table S1). The SAXS scattering profile can be completely superimposed on the theoretical profile calculated from the *Cr*PRK crystal structure (Fig. 3*A*), indicating that the solved crystal structure closely matches the structure of the protein in solution (Fig. 3*B*).

**Fig. 3.**
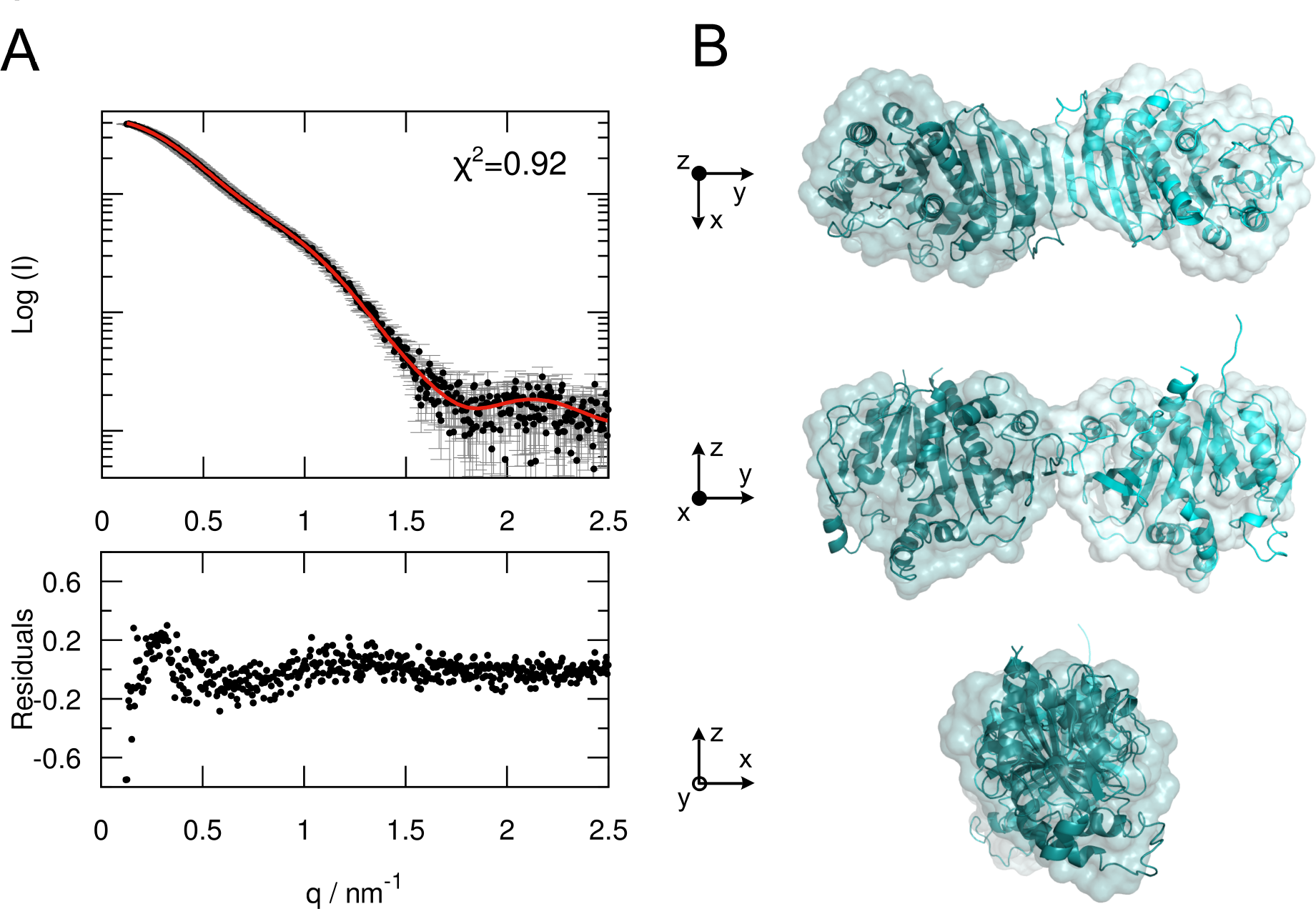
SAXS analysis of *Cr*PRK. (*A*) Superimposition of the experimental SAXS scattering profile (black) and the theoretical SAXS curve (red), calculated from the *Cr*PRK crystal structure (upper graph). The lower graph shows the residuals obtained by the difference between the experimental and calculated intensities. (*B*) Comparison between the ab initio model (surface representation) and the crystal structure (ribbon representation) of *Cr*PRK, showing an excellent agreement between the two models.

The active sites (one per monomer) are located in an elongated cavity at the edges of the dimer (Figs. 4*A* and S4*A*). In the *Cr*PRK structure, the active site cavity is marked by a sulfate ion derived from the crystallization medium that highlights the position occupied by the phosphate group of the substrate ribulose 5-phosphate (Ru5P) (Fig. 4*A* and B). The anion is stabilized by the side chains of Arg64, Arg67, and Tyr103, and by the main chain carbonyl group of His105 (Fig. S5). Except for Arg67, these residues are strictly conserved in PRKs (Fig. S6*A*). The catalytic role of the invariant Arg64 was also confirmed in *C. reinhardtii* (14). In addition, Arg64 was found to play a major role in the interaction of PRK with A_4_-GAPDH during the formation of the regulatory ternary complex (14). Consistent with this finding, Arg64 faces out at the far end of the dimer (Fig. 1 and Table S2). The active site also contains a pair of regulatory cysteines (Figs. 4*A* and *B* and S4*A* and *B*); Cys15 in *At*PRK and Cys16 in *Cr*PRK are located within the P-loop (Walker A) (15) and belong to the ATP-binding site (16). Additionally, Cys54 in *At*PRK and Cys55 in *Cr*PRK are located at the C-terminal end of strand β2 and compose the intermolecular mixed disulfide with TRX-f (6) (Figs. 2 and S2).

**Fig. 4.**
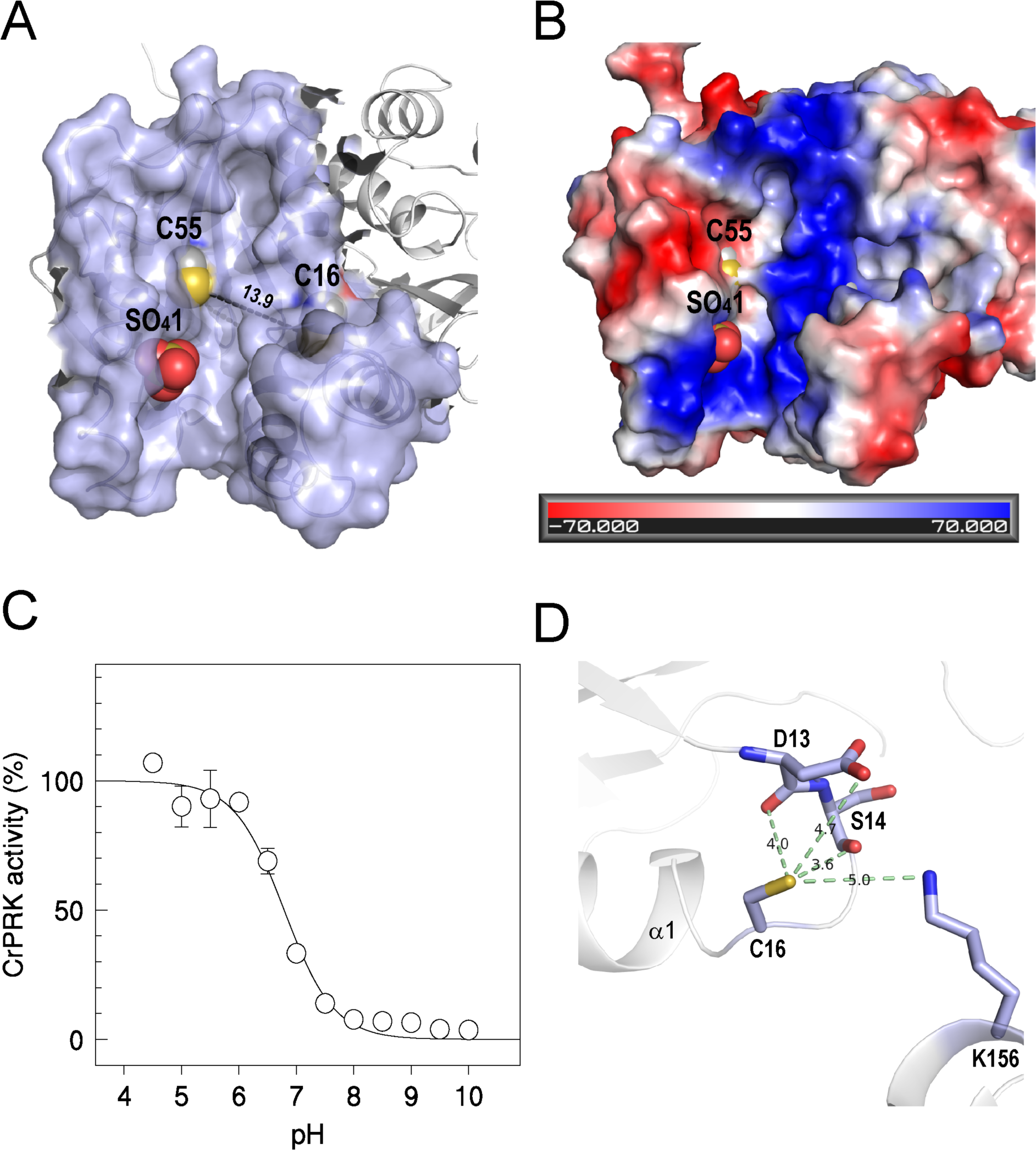
The active site and TRX-dependent regulation of *Cr*PRK. (*A*) The catalytic cavity is shown. The distance between the regulatory cysteines is greater than 13 Å. (*B*) The electrostatic surface potential of the catalytic cavity. The bottom of the catalytic cavity is marked by a positive potential. The negative potential observed in the left side of the cavity is thought to be involved in the correct positioning of TRX close to the regulatory cysteines. (*C*) The p*K*_a_ of Cys16 was determined by measuring the IAM-mediated inactivation as a function of pH. (*D*) The molecular environment of Cys16, considering a sphere of 5 Å centered on its thiol group. The hydrogen bonds between the thiol group and the neighboring residues, and the corresponding distances are shown.

### Redox properties of *Cr*PRK and *At*PRK

Despite the role that PRK plays in photosynthesis, putative PRK sequences have also been found in heterotrophic prokaryotes. Phylogenetic analyses performed on 69 PRK sequences, including 31 from nonphotosynthetic prokaryotes, clearly show a first evolutionary separation between Bacteria and Archaea. Cyanobacterial and eukaryotic PRKs emerged as the third clade from Archaea (Fig. S6*B*). Single PRK structures from the anoxygenic photosynthetic bacterium *Rhodobacter sphaeroides* (*Rs*PRK; PDB ID code 1A7J) (17) and the nonphotosynthetic Archaea *Methanospirillum hungatei* (*Mh*PRK; PDB ID code 5B3F; 18) have been reported. Both *Rs*PRK and *Mh*PRK sequences are shorter than plant PRKs with a low sequence identity (Fig. S7). All PRKs from bacteria (type I PRKs; 19), including phototrophic organisms, are octamers (~32-kDa subunits) that are allosterically activated by NADH and inhibited by AMP (20). Archaea, cyanobacteria, and eukaryotic (i.e., plant-type) PRKs are dimers (~40-kDa subunits; type II PRKs) (19) with no allosteric regulation (11, 21). While little is known about the regulation of archaeal PRKs, in oxygenic phototrophs from ancient cyanobacteria to modern flowering plants, it is well established that PRK forms a complex with A4-GAPDH and the regulatory protein CP12 at low NADP(H)/NAD(H) ratios (22, 23). Moreover, cyanobacterial PRKs are inhibited by AMP via the cyanobacteria-specific fusion protein CBS-CP12 (24), whereas plant-type PRKs are directly regulated by the TRX-dependent interconversion of specific dithiol-disulfide bridges (6). To date, no experimental evidence of the direct regulation of cyanobacterial PRKs by TRXs has been reported.

Regulatory cysteines Cys15/16 and Cys54/55 in *At*PRK and *Cr*PRK, respectively, protrude from the catalytic cavity (Figs. 4*A* and S4*A*) and are thus available to interact with TRX (Table S2). In addition, both PRKs share similar redox properties with comparable midpoint redox potentials (– 312 ± 3 mV for *Cr*PRK and –330 mV for *At*PRK) (23) and p*K*a values (6.79 ± 0.01 for *Cr*PRK and 6.95 ± 0.13 for *At*PRK) (Figs. 4*C* and S4*C*). These slightly acidic values are supported by the molecular environment around Cys15/16, as the thiol groups form hydrogen bonds with the main chain atoms of two nearby residues (i.e., Asp13 and Ser14 in *Cr*PRK and Asp12 and Ser13 in *At*PRK) and with the amino and carboxyl groups of Lys156 and Asp13 in *Cr*PRK or Lys151 in *At*PRK (Figs. 4*D* and S4*D*). All of these residues are strictly conserved in type I PRKs (Fig. S6*A*). Furthermore, the proximity of Cys15/16 to the positive N-terminal end of helix α1 contributes to a low p*K*a value (Figs. 4*D* and S4*D*).

The pH and temperature dependence of *At*PRK and *Cr*PRK are also comparable, although the algal enzyme has a wider range of activity at higher temperatures and lower pH values (Fig. S8). The pair of cysteines involved in TRX regulation is also present in the TRX-insensitive PRK in cyanobacteria (Fig. S6*A*). However, cyanobacterial PRKs lack an unstructured amino acid stretch (25)—here named clamp loop—which connects the regulatory cysteines whose thiol groups are separated by more than 13 Å, as found in plant-type PRKs (Figs. 4*A* and S4*A*). The clamp loop connects α1 and β2 (Figs. 1 and S7) and possibly confers the flexibility necessary to permit the conformational changes required for disulfide formation. This bond would thus allow the narrowing of the catalytic cavity and the inhibition of enzymatic activity (Fig. 5*A*). Similarly, a comparison of the chloroplast and redox-sensitive isoforms of FBPase (5) and AB-GAPDH (7) with their cytosolic and redox-insensitive counterparts reveals that the acquisition of TRX-mediated regulation is accompanied by the appearance of loops presumably required for the approach of the two cysteines involved in the formation of the regulatory disulfide bond.

**Fig. 5.**
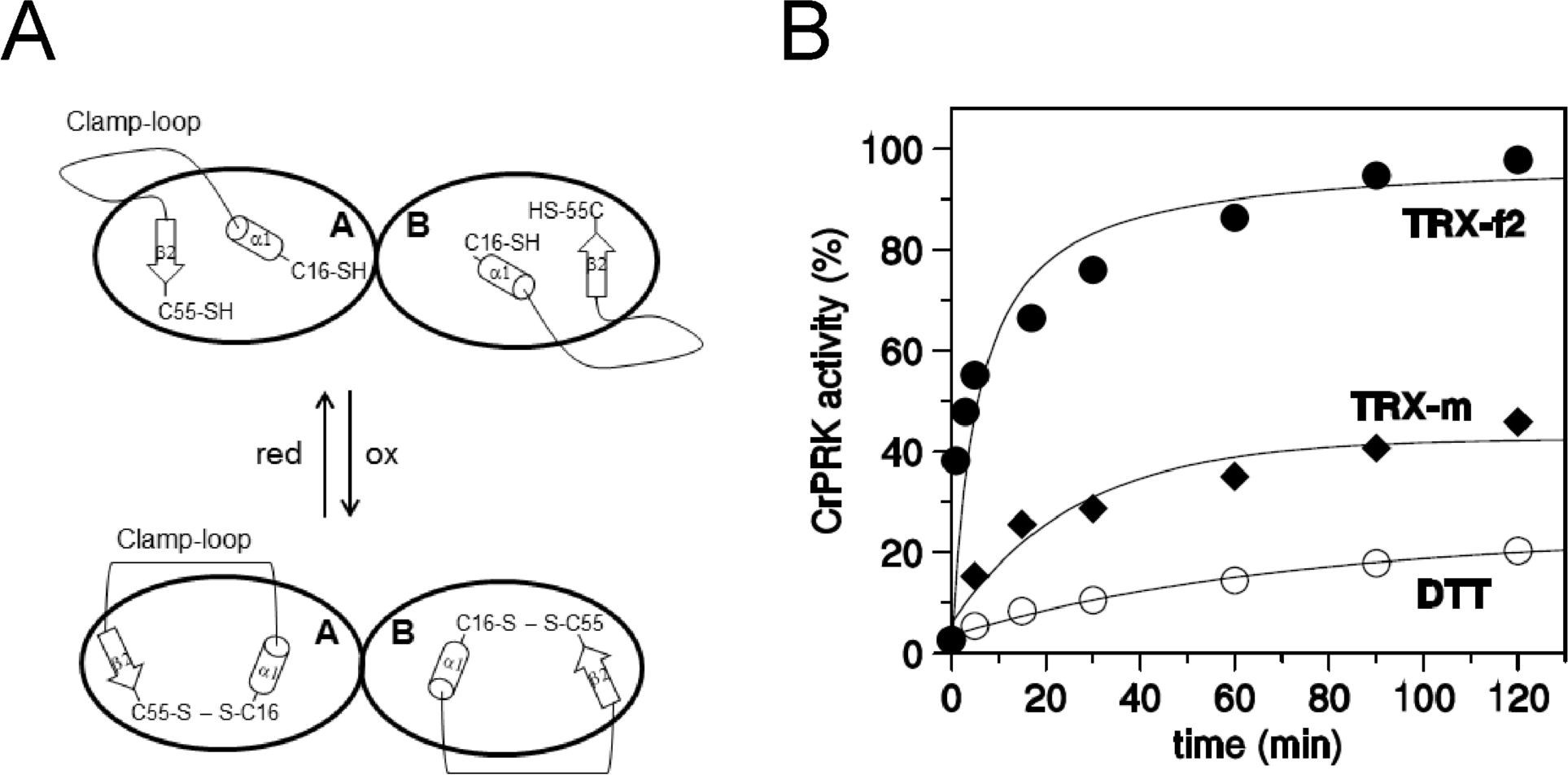
The clamp loop and TRX-dependent regulation of *Cr*PRK. (*A*) A schematic representation of the proposed role for the clamp loop and conformational changes occurring in the chloroplast PRK upon TRX binding. (*B*) Activation kinetics by plastidial *C. reinhardtii* TRX-f2 and -m, and DTT.

The electrostatic surface potential of the catalytic cavity reveals an elongated positive region on the bottom and a negative region on the side (Figs. 4*B* and S4*B*), the former suitable for binding the phosphate groups of substrates (i.e., ATP and Ru5P) and the latter relevant for the recognition and positioning of TRX, as proposed for FBPase (13, 26). Compared to TRX-m, TRX-f2 is more efficient in the reductive activation of both *At*PRK (27) and *Cr*PRK (Fig. 5*B*). Accordingly, the structural model of *At*TRX-f1 and the crystal structure of *Cr*TRX-f2 (28) show that the catalytic cysteines are surrounded by a large positive region, less extended in TRX-m (Fig. S9). This finding suggests that the early stages of pairing between TRX and PRK are mainly governed by electrostatic interactions between exposed regions of the two proteins (26).

Moreover, it has been reported that the oxidized state of target proteins is an essential feature for TRX target recognition, since disulfide formation does not confer any specific conformational change in TRXs structure (13). Indeed, the disulfide bond between the two distant regulatory cysteines introduces a significant conformational constraint that decreases the overall entropy of PRK. Therefore, a favorable entropic contribution is proposed to be the main driving force of the reduction of PRK by TRXs.

### Dimer interface and flexible regions

Dimer interfaces consist of large contact areas in octameric *Rs*PRK (1,667 Å^2^) (17) and dimeric *Mh*PRK (1,695 Å^2^; Fig. S10) (18), and a small area in eukaryotic PRK. The dimer interfaces in *Cr*PRK (545.6 Å^2^) and *At*PRK (560.3 Å^2^) are exclusively formed by β7, which is located in an antiparallel position to its partner (Figs. 1, 2, and S2), suggesting a less rigid structure. Since cyanobacterial and eukaryotic PRKs are found together with CP12, the greater flexibility of PRK could enable the formation of regulatory GAPDH/CP12/PRK complexes (9–11). Moreover, *At*PRK and *Cr*PRK are characterized by an increased number of exposed random coiled regions (Fig. 1 and Table S3), some of which show a poor or absent electron density, indicative of highly flexible and disordered regions (Fig. S11). Two such regions are common to algal and plant enzymes, whereas the presence of a third region is exclusive to *At*PRK.

The long loop between helices α5 and α6 (Fig. S11) contains the highly conserved catalytic Asp160 or Arg159 and Arg164 residues involved in Ru5P binding (29, 30). A second flexible region, corresponding to the loop between β7 and β8, contains Cys243 and Cys249 (Figs. 1 and S11). In both PRK structures, this portion is so flexible that the two thiol groups are far from each other in one monomer and closer to one another in the other; however, in contrast to *At*PRK, they are not oxidized in *Cr*PRK (Figs. 1 and S12). This result is in agreement with the observation that in *Cr*PRK, the disulfide bond between Cys243 and Cys249 only forms via an interaction with CP12 (31). This interaction is an essential step for the assembly of PRK and A4-GAPDH, leading to the regulatory complex GAPDH/CP12/PRK (32, 33).

## CONCLUDING REMARKS

TRXs are widespread regulatory proteins whose structure is strongly conserved throughout evolution. Fascinating experiments in paleobiochemistry conducted on seven “resurrected” TRXs dating back to the Precambrian era showed that only small differences in the length of helix α1 were found to occur over 4 Gyr of evolution (3). How evolution affected the TRX dependence of enzymes in the CB cycle is still a mystery. After studying the five CB cycle enzymes whose 3D structures have been solved or modeled (Fig S1), several strategies were adopted to establish their dependency on the same regulator. In fact, each of these five proteins are regulated by TRX in a different way. In FBPase and SBPase, the TRX-binding site is far from the conserved sugar-binding domain (5). In FBPase, upon reduction, the substrate-binding site adopts a catalytically competent conformation due to the loosening of secondary structures (4, 5). In AB-GAPDH, the redox-active cysteines are found in the highly flexibly CTE, which is characteristic of B-subunits, and when oxidized, the last residue in the CTE blocks the NAPDH-binding site (7). Similarly, a change in protein flexibility is believed to enable the regulation of *Chlamydomonas* PGK in which the formation of the regulatory disulfide appears to lower catalytic efficiency by decreasing the flexibility required to approach the substrates (8). Finally, we report that the regulatory pair of cysteines in PRK is embedded in the substrate-binding site and that oxidation can be achieved by the presence of the flexible clamp loop.

We conclude that a common strategy has allowed the acquisition of TRX dependence of the CB cycle enzymes during evolution—the introduction of flexible regions. This strategy is in sharp contrast to the evolutionary rigidity of the TRX structure.

## MATERIALS AND METHODS

### Protein Expression and Purification

Recombinant *At*PRK and *Cr*PRK were expressed in *E. coli* BL21(DE3) strain (New England Biolabs) harboring the pET-28 expression vector (Novagen) containing the coding sequence of the mature form of both enzymes (Fig. S7). *At*PRK was expressed and purified as previously described (33). The sequence coding for *Cr*PRK was cloned in frame with a 6xHis-Tag and a thrombin cleavage site at the 5’ end of the nucleotide sequence. *Cr*PRK was purified by Chelating Sepharose Fast Flow (GE Healthcare) column, following the manufacturer instruction. Immediately after elution, *Cr*PRK was desalted in 30 mM Tris-HCl, 1 mM EDTA, pH 7.9 by PD-10 Desalting Columns (GE Healthcare). The 6xHis-Tag was removed by overnight digestion with thrombin protease. Samples purity was checked by 12.5% SDS-PAGE and purified proteins were quantified by absorbance at 280 nm (Nanodrop; Thermo Scientific) using the following extinction coefficients: ε*At*PRK 31985 M^-1^ cm^-1^; ε*Cr*PRK 35995 M^-1^ cm^-1^ and molecular weights: *At*PRK 38784.23 g mol^-1^ and *Cr*PRK 40790.7 g mol^-1^. Proteins were stored at −80 °C.

When required, oxidized and reduced *At*PRK and *Cr*PRK were prepared by incubation at 25°C for 2-3 h in the presence of 40 mM trans-4,5-dihydroxy-1,2-dithiane (oxidized DTT) or 1,4-dithiothreitol (reduced DTT). Following incubation, samples were desalted in 30 mM Tris-HCl, 1 mM EDTA, pH 7.9 through NAP-5 column (GE Healthcare) and brought to the desired concentration either by dilution or by concentration through Amicon-Ultra device (Millipore; cut-off 10 kDa).

### Activity Assay and Determination of pH and Temperature Dependence

Phosphoribulokinase activity was measured spectrophotometrically (Cary 60 UV-Vis; Agilent) as previously described (34). The pH dependence of purified enzymes was determined in Britton-Robinson buffer for pH values ranging from 4.0 to 8.5. For pH ranging from 9 to 10, 100 mM Glycine buffer was used. Activities were measured on two independent protein batches and are expressed as percentage of the highest measured activity. Temperature dependence was evaluated on aliquots of *At*PRK and *Cr*PRK incubated for 20 min at temperatures ranging from 20°C to 60°C. Following incubation, the activity was assayed at 25°C. Activities were measured on two independent protein purifications and are expressed as percentage of the highest measured activity.

### Thioredoxin Specificity, Redox Titration

Thioredoxin specificity for activation of oxidized *Cr*PRK (2 µM) was measured in the presence of 0.2 mM reduced DTT and *C. reinhardtii* recombinant TRX-f2 and TRX-m (5 µM each), as previously described (27). At different incubation times, PRK activity was measured. To determine the midpoint redox potential of *Cr*PRK, three independent redox titrations were performed. Briefly, a mixture containing pure recombinant *Cr*PRK (12 µM), commercial *E. coli* thioredoxin (1 mg ml^-1^) and 20 mM DTT, at different reduced to oxidized ratios, were incubated for 3 h at 25°C before measuring PRK activity. The obtained curves were fitted by non-linear regression (CoStat, Co-Hort Software) to the Nernst equation.

### Determination of the p*K*_a_ Values of the Catalytic Cysteines

The determination of p*K*_a_ values for both *Cr*PRK and *At*PRK was performed as previously described (35). Briefly, PRK activity was measured on reduced PRK samples (2 µM) following 20 min of incubation at 25°C in the presence or absence of 0.2 mM iodoacetamide (IAM). Incubations were performed in different buffers within a 4.5-10 pH range. Aliquots of 10-20 µl were used to measure PRK activity in 1 ml of assay mixture. The residual activity was expressed as percentage of inhibition between IAM-treated and untreated samples, and expressed as a function of pH. The obtained curves were fitted by non-linear regression (CoStat, Co-Hort Software) to modified Henderson-Hasselbalch equation.

### Crystallization Data Collection and Structure Refinement

Reduced *Cr*PRK was concentrated at 10 mg ml^−1^ in 30 mM Tris-HCl, 1 mM EDTA, pH 7.9. Reduced *At*PRK was concentrated at 10 mg ml^−1^ in 25 mM potassium phosphate buffer solution, pH 7.5. They were crystallized by the vapor diffusion method at 293 K with sitting drop in 22% w/v PEG MME 5K, 0.1 M MES, pH 6.5, 0.2 M ammonium sulfate for *Cr*PRK, and hanging drop in 1.5 M sodium malonate, pH 5.0, for *At*PRK. The crystals obtained were analyzed by X-ray diffraction, and the structures were solved by molecular replacement. The refinement was performed with REFMAC 5.8.0135 (36) selecting 5% of reflections for R_free_ (for details, see SI Appendix, SI Materials and Methods).

### Small Angle X-ray Scattering Data Collection and Analysis

Small angle X-ray scattering (SAXS) data were collected at the BioSAXS beamline BM29 (37) at European Synchrotron Radiation Facility (ESRF, Grenoble) and the data collection parameters are reported in Table S4. A size-exclusion chromatography SEC-SAXS experiment was performed using a HPLC system (Shimadzu) directly connected to the measurement capillary. All the details about experimental procedures and data analysis are listed in SI Appendix, SI Materials and Methods.

### Modeling from SAXS Data

The experimental SAXS profile of the *Cr*PRK dimer was fitted with the theoretical profile calculated from the atomic coordinates using CRYSOL3 (38) with default parameters. In addition, low-resolution models of the CrPRK dimer based on the SAXS profile were built using the ab initio program GASBOR (39). All details are reported in SI Appendix, SI Materials and Methods.

### Data Availability

Atomic coordinates and structure factors have been deposited in the Protein Data Bank (PDB) under accession codes 6H7G and 6H7H for *Cr*PRK and *At*PRK, respectively. The SEC-SAXS data of *Cr*PRK have been deposited in SASBDB with accession code SASDDH9.

## ACKNOWLEDGEMENTS

We sincerely thank Prof. Bob B. Buchanan for offering comments and constructive suggestions on the content of the manuscript. We thank the Elettra and European Synchrotron Radiation Facility for allocation of X-ray diffraction and SAXS beam time (BAG proposal MX686 and MX1750) and the staff of beamlines XRD1 (Elettra) and BM29 (ESRF) for technical support. This work was supported by the Italian Minister of Education, Research and University (MIUR) grant FIRB2003 (P.T.); by University of Bologna grant FARB2012 (F.S., S.F., and M.Z.); by CNRS, Sorbonne Université, Agence Nationale de la Recherche Grant 17-CE05-0001; and CalvinDesign and LABEX DYNAMO ANR-LABX-011 (J.H., S.D.L., and P.C.). S.F. thanks the Consorzio Interuniversitario di Ricerca in Chimica dei Metalli nei Sistemi Biologici (CIRCMSB).

