## Supplemental material for "*Arabidopsis* and *Chlamydomonas* phosphoribulokinase crystal structures complete the redox structural proteome of the Calvin-Benson cycle"

**This PDF file includes:**

Supplementary text

Figs. S1 to S13

Tables S1 to S6

References for SI reference citations

#### Supplementary Information

#### SI Materials and Methods

**Crystallization and Data Collection**.

Aliquots of 2 µl for both proteins were mixed to an equal volume of reservoir, and the prepared drop was equilibrated against 900 (*Cr*PRK) and 750 (*At*PRK) µl of reservoir.

*Cr*PRK crystals were obtained in different conditions of the Extension Kit from Hampton Research (solutions 22: 12% w/v PEG 20K, 0.1 M MES, pH 6.5; 26: 30% w/v PEG MME 5K, 0.1 M MES, pH 6.5, 0.2 M ammonium sulfate; 30: 10% w/v PEG 6K, 5% v/v MPD, 0.1 M HEPES, pH 7.5) and Structure screen 1 from Molecular Dimension (MD1-01-CF, solutions 35: 30% w/v PEG 4K, 0.1 M Tris-HCl, pH 8.5, 0.2 M lithium sulfate; and 50: 15% w/v PEG 8K, 0.5 M lithium sulfate). The conditions were optimized and the best diffracting crystal grew in about 10 days, from 22% w/v PEG MME 5K, 0.1 M MES, pH 6.5, 0.2 M ammonium sulfate. *Cr*PRK crystal appeared as needle-like, in the majority of the cases forming a cluster. Aggregates were manually separated and the thicker individuals fished.

*At*PRK crystals showed a bipiramidal morphology and grew in three to five weeks from a reservoir solution containing 1.4 – 1.7 M sodium malonate, pH 5.0. The best diffracting crystal of *At*PRK was obtained in 1.5 M sodium malonate, pH 5.0.

Crystals were mounted from the crystallization drop into cryo-loops, briefly soaked in a cryo-protectant solution containing 30% w/v PEG MME and 20% v/v PEG 200 for *Cr*PRK and 1.7 M sodium malonate, pH 5.0, and 30% v/v glycerol for *At*PRK, then frozen in liquid nitrogen. Diffraction images were recorded at 100 K at the Elettra synchrotron radiation source (Trieste, beam line XRD1) for *Cr*PRK and at the European Synchrotron Radiation Facility (Grenoble, beam line ID14-4) for *At*PRK. Data collection parameters are reported in Table S4.

The data at a resolution of 2.6 Å for *Cr*PRK and 2.5 Å for *At*PRK were processed using XDS (1) and scaled with SCALA (2). The correct space group was determined with POINTLESS (2) and confirmed in the structure solution stage. Data collection statistics are reported in Table S5.

**Structure Solution and Refinement.**

*Cr*PRK structure was solved by molecular replacement using the program PHASER (3) from PHENIX (4) starting from the coordinates of PRK from *Methanospirillum hungatei* (PDB code 5BF3) (5) deprived of sulfate ions and water molecules. The protein chain was traced by Autobuilt from PHENIX (4) and Buccaneer (6) from CCP4 package. The refinement was performed with REFMAC 5.8.0135 (7) selecting 5% of reflections for $R_{free}$. The manual rebuilding was performed with Coot (8). The residual electron density map showed the position of a sulfate ion for each monomer coming from the crystallization solution, which was added to the model. Water molecules were automatically added and, after a visual inspection, confirmed in the model if the relative electron density value in the ($2F_o - F_c$) maps exceeded 0.19 $e^{-}Å^{-3}$ (1.0 σ) and if they fell into an appropriate hydrogen bonding environment. The last refinement cycle was performed with PHENIX (4). The structure of *Cr*PRK without sulfate ions and waters, was used as initial model to solve the *At*PRK structure by molecular replacement using MOLREP (9). The refinement was performed as described for *Cr*PRK. The refinement statistics are reported in Table S5. All structure figures were prepared using PyMOL (The PyMOL Molecular Graphics System, Schrödinger, LLC).

**Small Angle X-ray Scattering Data Collection.**

Data collection parameters are reported in Table S4. A Size Exclusion Chromatography SEC-SAXS experiment was performed using a HPLC system (Shimadzu) directly connected to the measurement capillary. A volume of 100 $\mu$l of reduced *Cr*PRK (6.1 mg $ml^{-1}$) was loaded onto a Superdex 200 10/300 GL column (GE Healthcare) pre-equilibrated in 50 mM Tris-HCl, 150 mM KCl, pH 7.5. The sample was eluted at a flow rate of 0.5 ml $min^{-1}$ and SAXS frames obtained by 1 s exposure were collected continuously. The automatic pipeline for SEC-SAXS data analysis implemented at BM29 was used to assess the quality of the collected data (10).

**Small Angle X-ray Scattering Data Analysis.**

Afterwards, a classification of the collected frames as buffer (0-14 ml) or protein frames (14-17.75 ml) was performed on the basis of the SAXS intensity trace (Fig. S13). Statistical test implemented in CorrMap (11) aided by visual inspection, was used to choose the superimposable buffer intensity profiles. The averaging of the buffer profiles, the subtraction of the averaged buffer intensity from the protein data and an automatic analysis of the subtracted protein profiles was performed with a Matlab script. The script used the tools of the ATSAS package (12) to automatically obtain from the subtracted intensity I(q): (i) the I(0) and the gyration radius ($R_g$) via the Guinier approximation (13) $I(q) = I(0)\cdot\exp[-(qR_g)^2/3]$; (ii) the pair-distance [p(r)] function, from which the maximum particle dimension ($D_{max}$) was estimated, in addition to an independent calculation of I(0) and $R_g$. Estimates of the MW were also determined both from the Porod invariant (14) as 0.6 times the Porod volume ($V_p$) for roughly globular particles (12) and by the invariant volume-of-correlation length ($V_c$), through a power-law relationship between $V_c$, $R_g$ and MW that has been parametrized (15). The protein frames giving constant $R_g$ values were scaled to the intensity of the elution maximum and averaged in order to obtain a single representative scattering profile with good signal to noise ratio, presented in the results and used for modelling.

**Modeling from SAXS Data**

The sequence and the homodimeric state of *Cr*PRK were given as inputs in GASBOR and a 2-fold symmetry was imposed in the calculations. A series of 10 models was generated. The similarity of the structures obtained by repeated calculations was checked by DAMAVER (16) in which the superposition is performed by the SUPCOMB code (17).

All programs used for SAXS data analysis and reconstruction, belong to the ATSAS package 2.7 (12). The graphical representations of the obtained three-dimensional models were built by using PyMOL (The PyMOL Molecular Graphics System, Schrödinger, LLC).

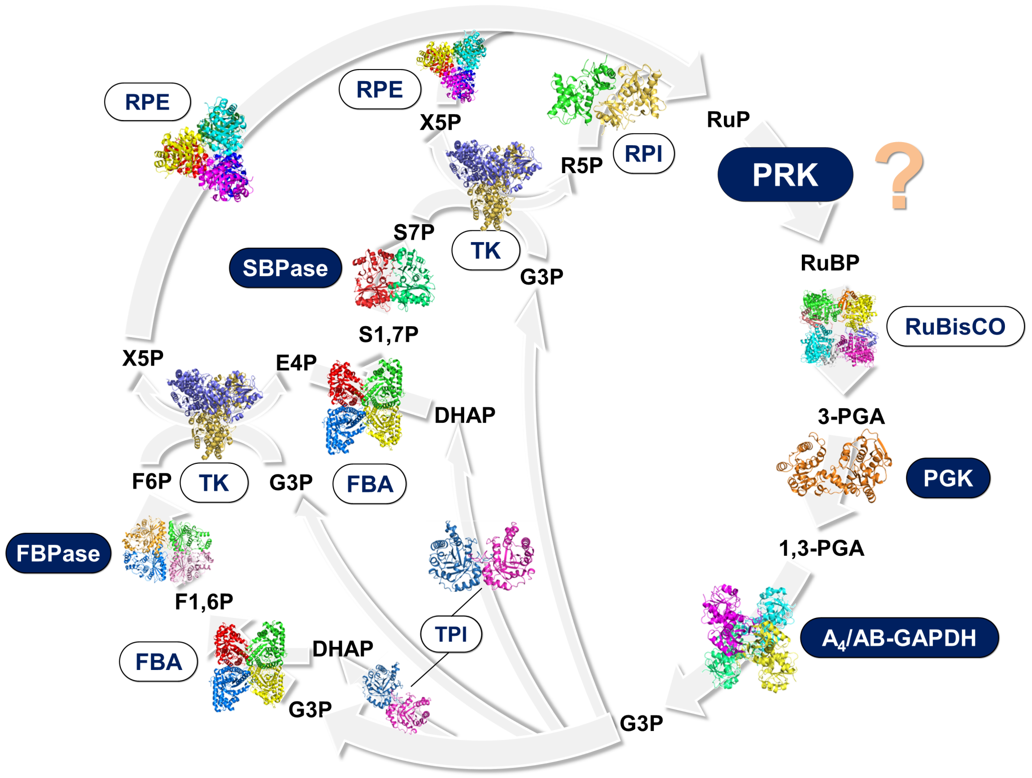


**Fig. S1. Schematic representation of the CB cycle.** The crystal structure of six chloroplast enzymes from different organisms: RuBisCO, ribulose-1,5-bisphosphate Carboxylase/Oxygenase from *Spinacia oleracia* (PDB ID code 1AUS) (18); GAPDH, glyceraldehyde-3-phosphatedehydrogenase from *Arabidopsis thaliana* (PDB ID code 3K2B) (19); TPI, triose phosphate isomerase from *Chlamydomonas reinhardtii* (PDB ID code 4MKN) (20); FBPase, fructose-1,6-bisphosphatase from *Pisum sativum* (PDB ID code 1DCU) (21); TK, transketolase from *Chlamydomonas reinhardtii* (PDB ID code 5ND5) (22); SBPase, sedoheptulose-1,7-bisphosphatase from *Physcomitrella patens* (PDB ID code 5IZ3) (23); RPE, ribulose-5-phosphate 3-epimerase from *Solanum tuberosa* (PDB ID code 1RPX) (24), is shown. For the remaining three enzymes, the crystal structure of the most homologous enzymes from non-photosynthetic organisms is reported: PGK, phosphoglycerate kinase from *Bacillus stearothermophilus* (PDB ID code 1PHP) (25) approximately 60% homologous to *Chlamydomonas reinhardtii* PGK1 (sequence accession number A8JC04); FBA, fructose-1,6-bisphosphatealdolase from *Toxoplasma gondii* (PDB ID code

5TJS) (26) approximately 55% homologous to *Arabidopsis thaliana* FBA1 (sequence accession number Q9SJU4); RPI, ribose-5-phosphateisomerase from *Toxoplasma gondii* (PDB ID code 4NML) approximately 50% homologous to *Spinacia oleracia* RPI (sequence accession number Q8RU73).

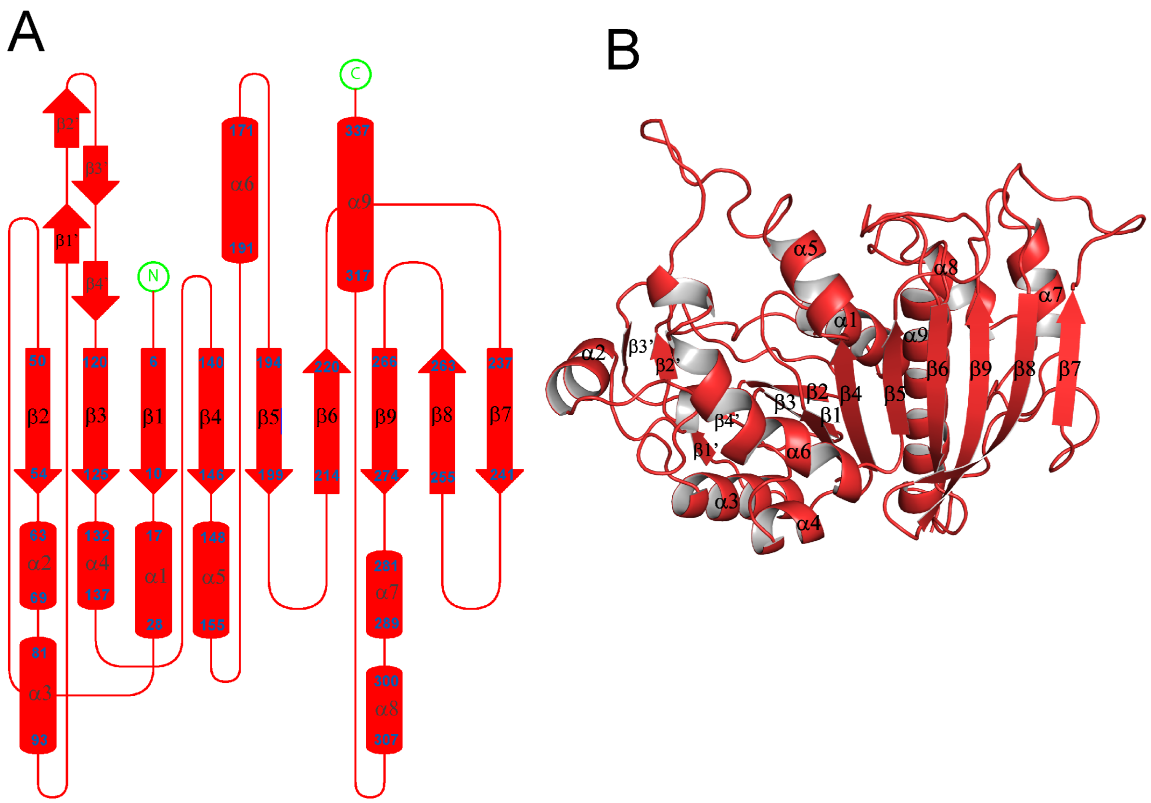


**Fig. S2. Crystal structure of *At*PRK's monomer.** (A) Topology diagram of *At*PRK. Similarly to *Cr*PRK (Fig. 2*A*), the monomer is composed by a mixed β-sheet of nine strands, by nine α-helices and four additional small β-strands indicated by β'. (B) Cartoon representation of the monomer structure of *At*PRK. Similarly to *Cr*PRK (Fig. 2*B*) the central β-sheet is sandwiched between helices α3, α4 and α6 and helices α1, α7, a8 and α9. The right end of the monomer consists of stand β7 involved in the dimer interface, while the four additional β-strands (β1' to β4') form the left external end of the dimer.

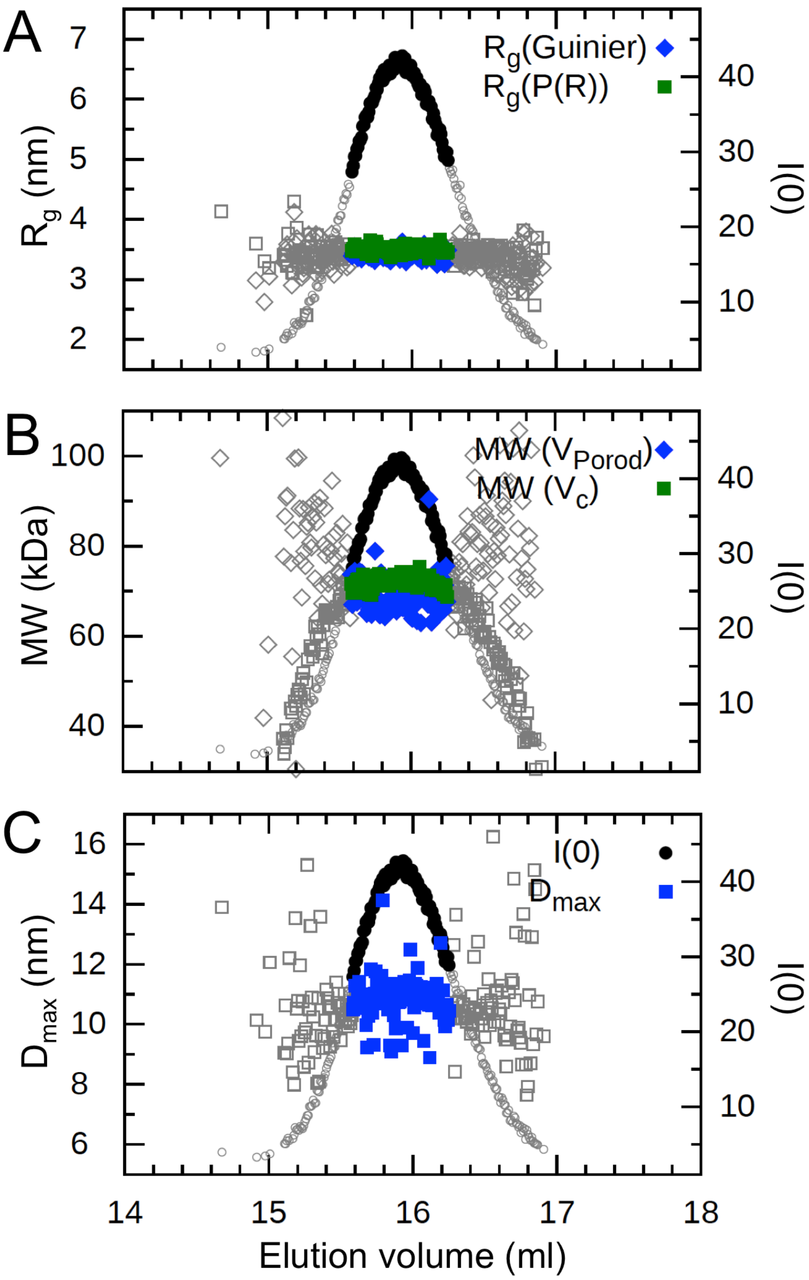


**Fig. S3. Parameters determined by SEC-SAXS analysis of *Cr*PRK.** (A) I(0) trace (dots) and $R_g$ determined by the Guinier approximation (diamonds) and $R_g$ calculated from the P(r) function (squares). (B) I(0) trace (dots) and MW estimated from the Porod volume (diamonds) and from the volume-of-correlation (squares). (C) I(0) trace (dots) and $D_{max}$ estimated from the P(r) function (squares). The frames used in the average to obtain the representative scattering profile, are highlighted in black compared to the grey neglected frames.

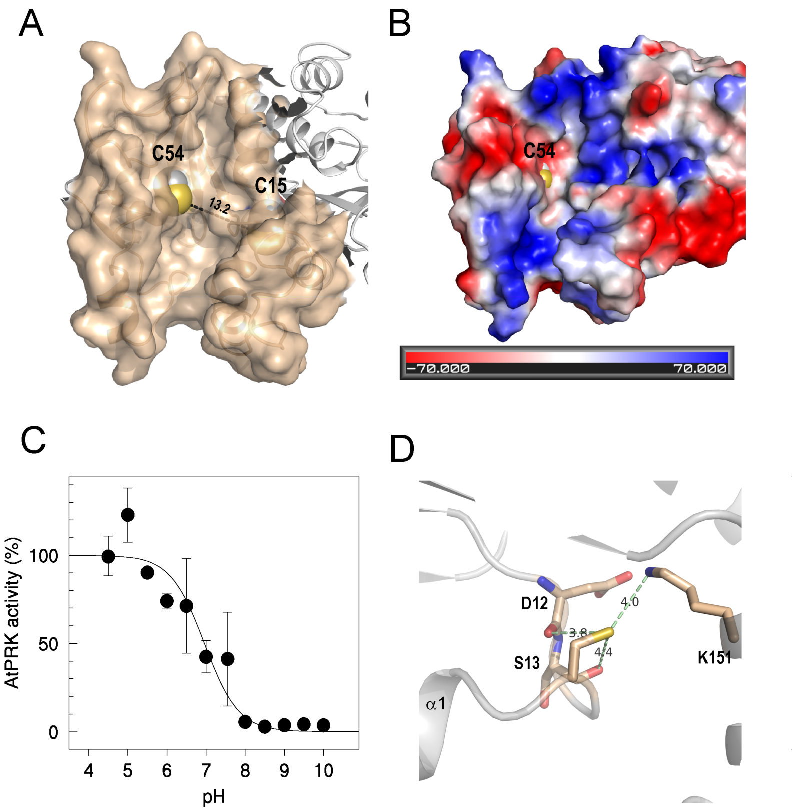


**Fig. S4. Active site and TRX-dependent regulation of AtPRK.** (A) The catalytic cavity is shown. The distance between the regulatory cysteines is higher than 13 Å. (B) Catalytic cavity electrostatic surface potential. The bottom of the catalytic cavity is marked by a positive potential. The negative potential region observed on the left side of the cavity, is suggested to be involved in the correct positioning of TRX close to regulatory cysteines. (C) The p$K_a$ of Cys15 was determined by measuring the IAM-mediated inactivation as a function of pH. (D) Molecular environment of Cys15 considering a sphere of 5 Å centered on its thiol group. The hydrogen bonds between the thiol group and the neighboring residues and the corresponding distances, are shown.

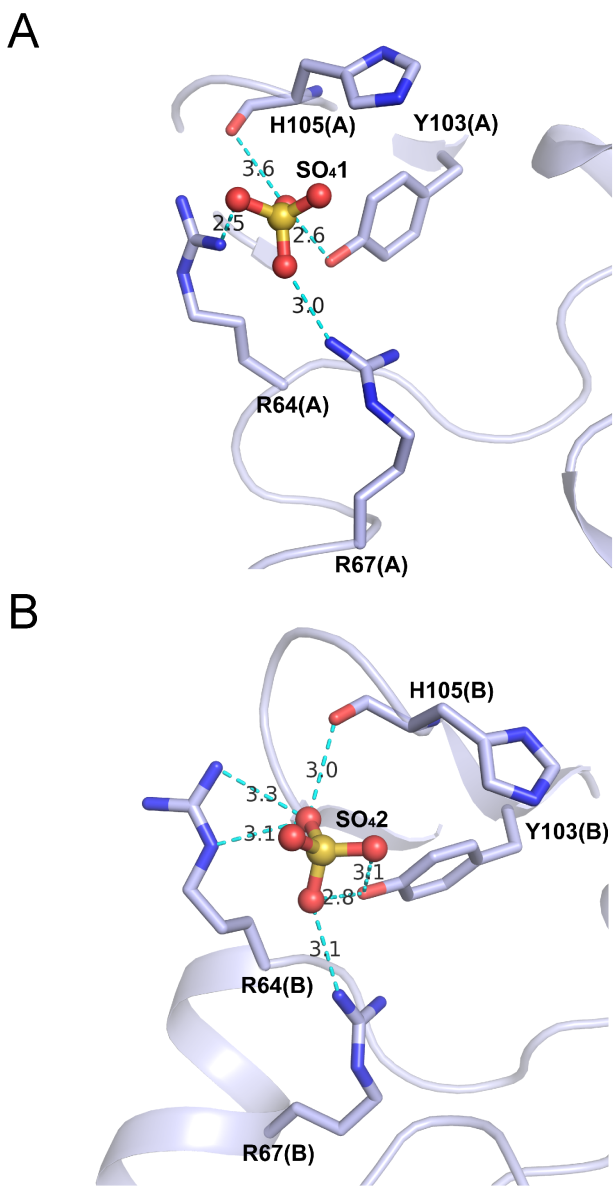


**Fig. S5. Binding site of the sulfate ion in *Cr*PRK**. The interactions between a sulfate ion coming from the crystallization solution, and protein residues in (A) subunit A and (B) subunit B, are represented. Salt-bridges are formed with Arg64 and Arg67, and hydrogen bonds with Tyr103 and His105.

A

```
                          10        20        30        40        50        60
                  =========+=========+=========+=========+=========+=========+
P. profundum      ------------------------------------------------------------
P. luminescens    ------------------------------------------------------------
E. amylovora      ------------------------------------------------------------
S. flexneri       ------------------------------------------------------------
S. medicae        ------------------------------------------------------------
R. meliloti       ------------------------------------------------------------
C. necator        ------------------------------------------------------------
R. rubrum         ------------------------------------------------------------
R. sphaeroides    ------------------------------------------------------------
A. cryptum        ------------------------------------------------------------
X. flavus         ------------------------------------------------------------
N. hamburgensis   ------------------------------------------------------------
N. vulgaris       ------------------------------------------------------------
R. palustris DX1  ------------------------------------------------------------
R. palustris Bis  ------------------------------------------------------------
M. capsulatus     ------------------------------------------------------------
A. ferrooxidans   ------------------------------------------------------------
C. M. oxyfera     ------------------------------------------------------------
T. denitrificans  ------------------------------------------------------------
A. vinosum        ------------------------------------------------------------
P. lunula         ---------------------------------MARSRSALFPALAALAALLLWAREAFL
L. polyedrum      ---------------------------MARSRSVVLPLAAAAVALVLLGHVAFVPPAPRP
B. natans         ------KTNVYAIASVVANVALVGVLLYTQTGEVGASVAASSRVHVAPRMVAPKFSMSGL
E. gracilis       -------------------------MMRYDQLIHEGPRQSSPSLLQLASVSATVFCAVFG
P. parvum         ------------------------------------------------MLSLSVGLSVAF
G. theta          -------------------------------------------MGTPLASRSSFASPVAL
T. pseudonana     ------------------------------------------------------------
O. sinensis       -------------------------------------MKLSLGIVALMAASASAFAPTAF
P. tricornutum    ---------------------------------------MKFAVFASLTATAAAFAPTAF
D. lutheri        ------------------------------------------------MRRSRVATPVQR
O. tauri          ------------------------------------------------------------
M. commoda        --------------------------------------------MASMTMSAACAQPLAA
P. sitchensis     ---------------------------------MACPAPTATSTPATVRSWNSQFCSNSP
P. patens         --------------------------------------MAAMACTSAASTALSTPRLAAV
S. oleracea       --------------------------------------MAVCTVYTIPTTTHLGSSFNQN
A. thaliana       --------------------------------------MAVSTIYSTQALNSTHFLTSSS
S. moellendorf.   --------------------------------------MAAACSSSGSSSRIGWFQASSS
T. aestivum       ------------------------------------------MAFCSPHTTTSLRSPCTT
P. trichocarpa    --------------------------------------MAVCTVYTTQSLNSTCSISTPT
M. crystallinum   --------------------------------------------------MAVSAYTVPT
P. sativum        ------------------------------------------------------------
C. variabilis     ---------------------------MQTAAAVQCKPVCRPMAARPHVARRPAQLAASA
C. reinhardtii    ------------------------------------------------------------
V. carteri        -----------------------------------------------MAFTMRAPAPRAT
G. sulphuraria    MELRDSLSTMGFVLPSTTSYVVYKSSSTIKSKKYNKRNYSSVSRVQVGGSKISSWSSNRL
G. kilaueensis    ------------------------------------------------------------
G. violaceus      ------------------------------------------------------------
T. elongatus      ------------------------------------------------------------
Synechocystis sp  ------------------------------------------------------------
M. aeruginosa     ------------------------------------------------------------
Cyanothece sp     ------------------------------------------------------------
S. elongatus      ------------------------------------------------------------
A. variabilis     ------------------------------------------------------------
N. spumigena      ------------------------------------------------------------
A. boonei         ------------------------------------------------------------
F. placidus       ------------------------------------------------------------
A. profundus      ------------------------------------------------------------
A. veneficus      ------------------------------------------------------------
M. harundinacea   ------------------------------------------------------------
M. concilii       ------------------------------------------------------------
M. thermophila    ------------------------------------------------------------
M. hungatei       ------------------------------------------------------------
Methanolinea sp   ------------------------------------------------------------
M. boonei         ------------------------------------------------------------
M. limicola       ------------------------------------------------------------
M. petrolearia    ------------------------------------------------------------
M. palustris      ------------------------------------------------------------
M. marisnigri     ------------------------------------------------------------
M. liminatans     ------------------------------------------------------------
Gblocks
Annotation
```

```
                          70        80        90       100       110       120
                  =========+=========+=========+=========+=========+=========+
P. profundum      ------------------------------------------------------------
P. luminescens    ------------------------------------------------------------
E. amylovora      ------------------------------------------------------------
S. flexneri       ------------------------------------------------------------
S. medicae        ------------------------------------------------------------
R. meliloti       ------------------------------------------------------------
C. necator        ------------------------------------------------------------
R. rubrum         ------------------------------------------------------------
R. sphaeroides    ------------------------------------------------------------
A. cryptum        ------------------------------------------------------------
X. flavus         ------------------------------------------------------------
N. hamburgensis   ------------------------------------------------------------
N. vulgaris       ------------------------------------------------------------
R. palustris DX1  ------------------------------------------------------------
R. palustris Bis  ------------------------------------------------------------
M. capsulatus     ------------------------------------------------------------
A. ferrooxidans   ------------------------------------------------------------
C. M. oxyfera     ------------------------------------------------------------
T. denitrificans  ------------------------------------------------------------
A. vinosum        ------------------------------------------------------------
P. lunula         LPRSSPQAPRASSARVPKVVALRAAADSQPTGLVWPSPEAEAKLREVDGIKLYPTHAWTE
L. polyedrum      RAARALTAMRGAANSVPTGLVWPSPEAEAKLREEDGIKMYPTHAWTDDMMPIVPATKE--
B. natans         RPAQREARVAAHGKGVVSTVALDAPVATSEDSMTWSSAKGNEVDAGMGQVHIEAAISNEV
E. gracilis       IGYTFSGGLVENYATTRPVQTQPAALILPKAVRYAGVSQGPQVESREARTALHAAATGTV
P. parvum         SPA--------------------------------------PSVVSQSRVAPRATVVE--
G. theta          -------------------------------------RVTNDKEMDVAIQQISMSLAS--
T. pseudonana     ------------------------MKFLVASLIASASAFSTINTPTLRGNSALAALKD--
O. sinensis       -----------------------------------MPANTLRAAAPASPSALNMALKE--
P. tricornutum    VPS-------------------------------------NLRGVAPSASSLNMALKE--
D. lutheri        VPQ--------------------------------------------TACTVLMATKT--
O. tauri          ---------------------MSASLGFSTSVRAAPVRATRDAVKPRATRARVVTTAK--
M. commoda        RKFQGS--------------------------------RVSGKSVVRAAKRNVTVKAE--
P. sitchensis     LAS-----------------------------------FSLAHTTRRPRRALVVCSVG--
P. patens         KPARSARPVH--------LTSAFHGQSVASVSQVAGFESSGVKYRSAGRRAVVVCKAA--
S. oleracea       NKQVFFNYKR----------------------------SSSSNNTLFTTRPSYVITCS--
A. thaliana       SSKQVF----------------------------------LYRRQPQTNRRFNTLITC--
S. moellendorf.   SSNIGILQHHPWRGSHASSLFSFPKLGARIGSSSGGSGSSGTSSSNRVRVLVCCAAGG--
T. aestivum       IPNSGFRQNQ----------------------VIFFTTRSSRRSNTRHGARTFQVSCA--
P. trichocarpa    KTHLGFNQRH-------------------------VVFYSTNKKTTKRASSAVITCSA--
M. crystallinum   TSHLGFNQKK---------------------QLFFCNKSAYKRVSFSSRPCVITCLAG--
P. sativum        --------------------------------------------------------AG--
C. variabilis     APIARPTFSS-------------------------TLNQRTLKAGRVASRVVVVKAEG--
C. reinhardtii    --------------------------MAFTMRAPAPRATAQSRVTANRARRSLVVRAD--
V. carteri        --------------------------------------AQSRVTASRASRRVLVVKAQ--
G. sulphuraria    LFQGKEINTTRKTTTKYWIITVSSQTALEEFVNCSGAKGVHESAGISRSKSKVLRNKG--
G. kilaueensis    ------------------------------------------------------MVSK--
G. violaceus      ------------------------------------------------------MVST--
T. elongatus      ------------------------------------------------------MSSK--
Synechocystis sp  ------------------------------------------------------MTTQ--
M. aeruginosa     ------------------------------------------------------MANK--
Cyanothece sp     ------------------------------------------------------MTTQ--
S. elongatus      -------------------------------------------------------MSK--
A. variabilis     ------------------------------------------------------MTTK--
N. spumigena      ------------------------------------------------------MTTK--
A. boonei         -----------------------------------------------MLGEFRRRLEE--
F. placidus       -----------------------------------------------------MILEK--
A. profundus      --------------------------------------------------MLKEKLIK--
A. veneficus      -----------------------------------------------MTSNLKERLKE--
M. harundinacea   ---------------------------------------------------MAERLKG--
M. concilii       ------------------------------------------------MRSLKDRIRE--
M. thermophila    -------------------------------------------------MRLLEKIRE--
M. hungatei       ---------------------------------------------MSQPENFREVIRH--
Methanolinea sp   ---------------------------------------------MTSKPGFKEIIKS--
M. boonei         ---------------------------------------------MPRTPPFKEIIAR--
M. limicola       -----------------MSCIDEYLSESGKCRNRFSGDDEYCFSSSGKGINLKKIIEN--
M. petrolearia    ----------------------------------------MDYSHETNLKNFRHAVDS--
M. palustris      -----------------------------MMQTEGKTGEKPDLCPGTGGLNFKDRIAS--
M. marisnigri     ----------------------------------------------MPPSDFKRVIAE--
M. liminatans     ----------------------------------------------MDTPVFRDLISG--
Gblocks
Annotation
```

```
                      130       140       150       160       170       180
                =========+=========+=========+=========+=========+=========+
P. profundum      -----------MSAKHPIIAVTGSSGAGTTTTSEAFRKMFNMM-----------------
P. luminescens    -----------MSAKHPIIAITGSSGAGTTTTSLAFRKIFQQL-----------------
E. amylovora      -----------MSTQHPVIAVTGSSGAGTTTTSLAFRKIFQQL-----------------
S. flexneri       -----------MSAKHPVIAVTGSSGAGTTTTSLAFRKIFAQL-----------------
S. medicae        -----------MSAKYPIISITGSSGAGTTTVKDTFEKIFKRE-----------------
R. meliloti       -----------MSAKFPIISITGSSGAGTTTVKDTFEKIFKRE-----------------
C. necator        -----------MSERYPIIAITGSSGAGTTSVTRTFENIFRRE-----------------
R. rubrum         -----------MSVKHPIIAITGSSGAGTTSVTRTFEQIFRRE-----------------
R. sphaeroides    -----------MSKKHPIISVTGSSGAGTSTVKHTFDQIFRRE-----------------
A. cryptum        -----------MSQRHPIISVTGSSGAGTTSVRNVFEQIFRRE-----------------
X. flavus         -----------MSIKHPIIVVTGSSGAGTTSVKRTFEQIFYRE-----------------
N. hamburgensis   -----------MSRKYPIISITGSSGAGTTSVKRTFEQIFRRE-----------------
N. vulgaris       -----------MLRKHPIISITGSSGAGTTSVKRTFEQIFRRE-----------------
R. palustris DX1 -----------MSRKHPIISITGSSGAGTTSVKKTFEQIFRRE-----------------
R. palustris Bis -----------MSRKHPIISITGSSGAGTTSVKRTFEQIFRRE-----------------
M. capsulatus     -----------MSKKHPIIAITGSSGAGTTTVKCAFEHIFFRL-----------------
A. ferrooxidans   -----------MSKKHPVIAVTGSSGAGTTTVKHAFHDIFRRL-----------------
C. M. oxyfera     -----------MSKKHPVIAVTGSSGAGTTFVKRAFENIFRRE-----------------
T. denitrificans  -----------MSKKHPVIAVTGSSGAGTTTVKRAFEHIFFRE-----------------
A. vinosum        -----------MSKKHPIVAVTGSSGAGTTTVKRAFEHIFHRD-----------------
P. lunula         EMTPILPATKQLDNTPVIIGVAADSGCGKSTFLRRILGALGTEVTP------------GH
L. polyedrum      ------------GVSPVVIGVAADSGCGKSTFLRRILGALGTEVTP------------GH
B. natans         SPVMAQKN----IQRPVIFGVAADSGCGKSTFLRRVNAIFGTTTSK------------AH
E. gracilis       NRDSTLQRPKVDPKKTVLIGVAADSGCGKSTFMRRLTGIFGGGKPTPLGGGFGTGGWETN
P. parvum         ------------MVNPVVVGVAADSGCGKSTFMRRLTSIFGGECKLLDIGRE------TN
G. theta          ------------GQKPVVIGLAADSGCGKSTFMRRVTACFGGACKLNPIGRE------TN
T. pseudonana     ------------GEVPIVIGLAADSGCGKSTFMRRVTSTFGGETCGPLGGGFGNGGWETN
O. sinensis       ------------GEKPIVIGVAADSGCGKSTFMRRLTSIFGGEGVGPLGGGFDNGGWETN
P. tricornutum    ------------GQTPIIIGVAADSGCGKSTFMRRLTNIFGGDVVGPLGGGFDKGSWETN
D. lutheri        ------------GVKAFVVGVAADSGCGKSTFMRRLTNIFGGKSALLDIGRE------TN
O. tauri          ------------RDGPVIIGLAADSGCGKSTFMRRMTSLFGGKATPPEGGNPD-----SN
M. commoda        ------------RDGPVLIGLAADSGCGKSTFMRRMTSLFGGKATPPEGGNPD-----SN
P. sitchensis     ------------PEKTVVIGLAADSGCGKSTFMRRLTSVFGGAAEPPKGGNPD-----SN
P. patens         ------------DGQTVVIGLAADSGCGKSTFMRRLTSVFGGAASPPKGGNPD-----SN
S. oleracea       ------------QQQTIVIGLAADSGCGKSTFMRRLTSVFGGAAEPPKGGNPD-----SN
A. thaliana       ------------AQETIVIGLAADSGCGKSTFMRRLTSVFGGAAKPPKGGNPD-----SN
S. moellendorf.   ------------DGKTIVIGLAADSGCGKSTFMRRLTSVFGGAASPPKGGNPD-----SN
T. aestivum       ------------VEQPIVIGLAADSGCGKSTFMRRLTSVFGGAAEPPKGGNPD-----SN
P. trichocarpa    ------------DTQTVVIGLAADSGCGKSTFMRRLTSVFGGAAEPPRGGNPD-----SN
M. crystallinum   ------------DSQTIVIGLAADSGCGKSTFMRRLTSVFGGAAEPPRGGNPD-----SN
P. sativum        ------------DSQTIVIGLAADSGCGKSTFMRRLTSVFGGAAEPPKGGNPD-----SN
C. variabilis     ------------GDKIVVIGLAADSGCGKSTFMRRVTGIFGGEPKPPSGGNPD-----SN
C. reinhardtii    ------------KDKTVVIGLAADSGCGKSTFMRRMTSIFGGVPKPPAGGNPD-----SN
V. carteri        ------------KDKTVVIGLAADSGCGKSTFMRRMTSIFGGVPKPPAGGNPD-----SN
G. sulphuraria    ------------IERPVIIGVAADSGCGKSTFLRRVNEIFGTKVSQ------------SH
G. kilaueensis    ------------ADRVVLIGVGGDSGCGKSTFLRRLKDLFSK------------------
G. violaceus      ------------LDRVVLIGVGGDSGCGKSTFLRRLKDLFSK------------------
T. elongatus      ------------PDRVVLIGVAGDSGCGKSTFLRRLADLFGE------------------
Synechocystis sp ------------LDRVVLIGVAGDSGCGKSTFLRRLTDLFGE------------------
M. aeruginosa     ------------PERVVLIGVAGDSGCGKSTFLRRLTDLFGA------------------
Cyanothece sp     ------------ADRVVLIGVAGDSGCGKSTFLRRLTDLFGE------------------
S. elongatus      ------------PDRVVLIGVAGDSGCGKSTFLNRLADLFGT------------------
A. variabilis     ------------PERVVLIGVAGDSGCGKSTFLRRLIDLFGE------------------
N. spumigena      ------------PERVVLIGVAGDSGCGKSTFLRRLIDLFGE------------------
A. boonei         ------------YEGSLIIGVAGDSGSGKSTFTKSIINLLGK------------------
F. placidus       ------------LKEPFLIGVAGDSGSGKTTFAESVRKIFAGN-----------------
A. profundus      ------------SGKVFIIGIAGDSGSGKTTFANGIKRMFGD------------------
A. veneficus      ------------SGKTFLVGIAGDSGSGKTTFANAIRNLLGN------------------
M. harundinacea   ------------SGRVFVVAVAGDSGSGKTTLSRGIRRLLGE------------------
M. concilii       ------------SGRVFVFGIAGDSGSGKTTISRGIRRILGE------------------
M. thermophila    ------------SGRVFVVAVAGDSGSGKTTFTRGIRRLLGE------------------
M. hungatei       ------------SPLVYLIGVAGDSGSGKSTFTRAISDIFGE------------------
Methanolinea sp   ------------SPGRFAIGVAGDSGSGKTTFTDSIRHLFGD------------------
M. boonei         ------------SPLVFVIGVAGDSGSGKTTFTNAIREIFGP------------------
M. limicola       ------------SGLIFVIGVSGDSGSGKTTFTDAIREIFGP------------------
M. petrolearia    ------------SESTFIIGVAGDSGSGKTTFTRAIRSIFGN------------------
M. palustris      ------------SPCVFTIGVAGDSGSGKTTFTQSIRDIFGE------------------
M. marisnigri     ------------SPYVFVIGVAGDSGSGKTTFTRAIREIFGD------------------
M. liminatans     ------------SNSVFVIGVAGDSGSGKTTFTRAIREIVGA------------------
Gblocks
Annotation                           |  *  |||
```

Walker A (P-loop) [contains C16]    Clamp loop

642

643

```
                             190       200       210       220       230       240
                  =========+=========+=========+=========+=========+=========+
P. profundum      ----NINASWLEGDSFHRYTRPEMDVEIRKAKEQGR-HISYFGPEANDFPQLEKFFRQYG
P. luminescens    ----DISAAQIEGDSFHRYTRPEMDAAIRKAKEQGR-HISYFGPEANDFGMLEKTMIDYG
E. amylovora      ----GLHAAEIEGDSFHRFTRPEMDMAIRKARDMGK-HVSYFGPEANDFALLERTFSEYG
S. flexneri       ----NLHAAEVEGDSFHRYTRPEMDMAIRKARDAGR-HISYFGPEANDFGLLEQTFIEYG
S. medicae        ----NISASFIEGDAFHRYDRETMRSKIAEEKARGV-DFTHFSAEANELEILESVFAEYG
R. meliloti       ----NISASFIEGDAFHRFDRETMRSKIAEEKARGV-DFTHFSAEANELEILESVFAEYG
C. necator        ----GVKSVVIEGDSFHRYDRAEMKVKMAEAERTGNMNFSHFGEENNLFGELENLFRSYA
R. rubrum         ----GVNAAVVEGDSFHRNDRKAMKIAMAEAQKAGNANFSHFGPEANLFEELETLFRTYG
R. sphaeroides    ----GVKAVSIEGDAFHRFNRADMKAELDRRYAAGDATFSHFSYEANELKELERVFREYG
A. cryptum        ----KITAAHIEGDAFHRYDRAEMKTKMAEAEAAGNRHFSHFSPETNLLAELAATFESYA
X. flavus         ----KVKAAFVEGDSFHRYDRYEMRELMAAEAAKGNKHFSHFSPETNRLDDLAQLFKDYG
N. hamburgensis   ----NVVAAYIEGDAFHRYNRADMRTRMAEESDRGNKHFSHFSPETNLFDELEAVFRSYS
N. vulgaris       ----NVVAAYIEGDAFHRYNRAEMRTRMAEESDKGNKHFSHFSPETNLFAELEGVFRSYG
R. palustris DX1  ----NVNAAFIEGDAFHRYNRVDMRNKMAEEAERGNRHFSHFSPETNLFEELEQTFKSYA
R. palustris Bis  ----NVNAAYIEGDAFHRYNRVDMRTRMAEEAEKGNKTFSHFSPDTNLFEELETTFRDYS
M. capsulatus     ----GLKPLVIEGDSFHRYDRVEMRAQIDKARREG-RHFSHFSIEANILDELENVFRHYG
A. ferrooxidans   ----KIDPVVIEGDSFHRYNRNEMREAIAKAAADG-KTISHFGPEGNDFEALERLFREYG
C. M. oxyfera     ----EITSAIIEGDSFHSVTRAQFKERSAVEH-----NFSHFGPEANDFHALEALFKSYG
T. denitrificans  ----KINAAVIEGDSFHSLARVEFKEAVKKAEAEGNFSFSHFGPQANHFDKLAELFKTYG
A. vinosum        ----NISAAVIEGDSFHSYDRATMKAEMAAFAKRGE-SLSHFGPVANRFDLLEELFREYG
P. lunula         TAVGDMMTVICLDD-YHTNDRAGRKA----------TGLTALDARENDFALMGSQIEALK
L. polyedrum      TAIGDMMTVICLDD-YHTNDRAGRKA----------TGLTALDAKENDFALMGVQIEALK
B. natans         TPTGDLITTICLDD-FHTLDRTGRAD----------TGISALDVRANNFALMADQLKALK
E. gracilis       TLVSDKTTVMCLDD-YHLNDRAGRKV----------TGLTALDQRENNFDLMFEQMSSLK
P. parvum         TLVSDMTTVICLDD-YHKWDRTGRKSNPEWPN-----GITALHEACQDWDKMAADVLDLK
G. theta          TLISDMTTVICLDD-YHLNDRQGRKK----------TGLTALDPRENNFDLMYEQVKALK
T. pseudonana     TLVSDMATVICLDD-YHLNDREGRKV----------SGLTALNTAEQKFDLMFEHVKALK
O. sinensis       SLVSDLTTVLCLDD-YHLNDRNGRKV----------TQRTALDPLENNFDLMYEQIAALK
P. tricornutum    TLVSDLTTVICLDD-YHLNDRAGRKV----------TMRTALDPEENNFDLMYEQVKALK
D. lutheri        TLVSDKTTVICLDD-YHLYDRKGRSA----------NKITALHKDCQKWDLMAEQVAAIK
O. tauri          TLISETTTVLCLDD-YHLNDRAGRKT----------SGLTALNLKEQNFDLMYDQVKALK
M. commoda        TLISDSTTVLCLDD-YHLNDRNGRKE----------SGLTALNLKEQNFDLMYEQTKALM
P. sitchensis     TLISDTTTVICLDD-YHSLDRYGRKD----------KGVTALDPRANNFDLMYEQVKALK
P. patens         TLISDTTTVICLDD-YHSLDRYGRKE----------KAVTALDPKANNFDLMYEQVKALK
S. oleracea       TLISDTTTVICLDD-FHSLDRNGRKV----------EKVTALDPKANDFDLMYEQVKALK
A. thaliana       TLISDTTTVICLDD-YHSLDRYGRKE----------QKVTALDPRANDFDLMYEQVKALK
S. moellendorf.   TLISDTTTVICLDD-YHSLDRTGRKE----------KGVTALDPKANNFDLMYEQVKALK
T. aestivum       TLISDTTTVICLDD-YHSLDRTGRKE----------KGVTALDPKANDFDLMYEQVKAIK
P. trichocarpa    TLISDTTTVICLDD-YHSLDRTGRKE----------KGVTALDPRANNFDLMYEQVKAIK
M. crystallinum   TLISDTTTVICLDD-YHSLDRTGRKE----------KGVTALDPRANDFDLMYEQVKALK
P. sativum        TLISDTTTVICLDD-YHSLDRTGRKE----------KGVTALDPKANDFDLMYEQVKAIK
C. variabilis     TLISDMTTVICLDD-YHSLDRNGRKE----------AGVTALAPESQNFDLMYEQVKALK
C. reinhardtii    TLISDMTTVICLDD-YHCLDRNGRKV----------KGVTALAPEAQNFDLMYNQVKALK
V. carteri        TLISDMTTVICLDD-YHCLDRNGRKV----------KGVTALAPEAQNFDLMYNQVKALK
G. sulphuraria    TPQGELVTVICLDD-FHTLDRKGRAE----------KKVTALNPEANNFELMYQQIAALK
G. kilaueensis    ----ELVTVICLDD-YHSLDRKQRKE----------TGITALDPRANNFDLMAEQAAALK
G. violaceus      ----ELVTVICLDD-YHSLDRKQRKE----------TGITALDPRANNFDLMAEQATALK
T. elongatus      ----DFMTVICLDD-YHSLDRKQRKE----------MGITALDPRANNFDLMYEQIKALK
Synechocystis sp  ----EFMTVICLDD-YHSLDRQGRKA----------AGVTALDPRANNFDLMYEQIKTLK
M. aeruginosa     ----EFMTVICLDD-YHCLDRKQRKE----------VGVTALNPKANNFDLMYEQIKALK
Cyanothece sp     ----EFMTVICLDD-YHSLDRQGRKK----------AGVTALNPQANNFDLMYEQIKALK
S. elongatus      ----ELMTVICLDD-YHSLDRKGRKE----------AGVTALDPRANNFDLMYEQVKALK
A. variabilis     ----EFMTVICLDD-YHSLDRKQRKE----------TGITALDPRANNFDLMYEQIKALK
N. spumigena      ----EFMTVICLDD-YHSLDRKQRKE----------TGITALDPRANNFDLMYEQIKALK
A. boonei         ----DLVSSFSLDD-YHTEDRETRKK----------TGHLPLDPKINNLKLAAEHLSALQ
F. placidus       ------VATITLDD-YHKYGRKERKK----------LGITPLNPEMTNLDLFREHLKLIK
A. profundus      ----DIVSHITLDD-YHVYDREMRER----------LGITPLHPSANNLKLVETHLYLLK
A. veneficus      ----DIVSSITLDD-YHVYDRQTRWK----------LNITPLHPEANNLKLVETHLMLLK
M. harundinacea   ----EAVSTFSMDD-YHSLDREERRV----------RGMTPLNPEANRLDLLAEHLRSLR
M. concilii       ----DMVATFSMDD-YHSLDRRQRKE----------RNITPLHPDANQFDLLAEHLEALR
M. thermophila    ----DVVSTFSMDD-YHSLDRRQRKT----------LGITPLRPEANRFDLLAEHLAMLR
M. hungatei       ----ELVSSITVDD-YHLYDRKTRSE----------MGITPLLHTANNLKLLEENLMDLK
Methanolinea sp   ----SMVATITLDD-YHIYNRTQRDE----------IQITPLHPDANNLAGLERDLRLLK
M. boonei         ----DLVATITLDD-YHTYDREERHH----------LDITPLHPDANNLSGLEADVAHLK
M. limicola       ----SFVSTITLDD-YHILDREERKE----------KNITPLHPDANNFSLLEEHLSDLK
M. petrolearia    ----NMVTTISLDD-YHVLDRDERKR----------TGITPLSPEANNFSLLEEHVALLK
M. palustris      ----DLVTTITLDD-YHLYDREERKV----------RHITPLNPEANRLDQLEHDLVELT
M. marisnigri     ----DLVSTITIDD-YHRYDRQERKV----------LGITPLVPEANRFDLLEEHLAELK
M. liminatans     ----DLVATITLDD-YHRYDRKERKD----------LGITPLVPEANNLDLLADHIRALK
Gblocks
Annotation                  *  +  #   #
                          C55
```

```
                        250       260       270       280       290       300
               =========+=========+=========+=========+=========+=========+
P. profundum      DDGSGQFRRYLHTFDEAVP-----------YNQMPGSFTPWQKLPENTDVLFYEGLHGGV
P. luminescens    ETGEGRRRKYLHTYDDAVP-----------YNQLPGTFTPWESLPKQTDVLFYEGLHGGV
E. amylovora      RSGKGKSRKYLHTYDEAVP-----------WNQVPGTFTPWQPLAEPTDVLFYEGLHGGV
S. flexneri       QSGKGKSRKYLHTYDEAVP-----------WNQVPGTFTPWQPLPEPTDVLFYEGLHGGV
S. medicae        RRGVGRTRHYVHDDAEAVK-----------FGSDPGTFTDWEEFR-DSDLLFYEGLHGCA
R. meliloti       RRGVGRTRHYVHDEAEAAK-----------FGSDPGTFTDWEEFR-DSDLLFYEGLHGCA
C. necator        ETGTGMHRHYLHSPEEAAP-----------FGQEPGTFTQWEPLPADTDLLFYEGLHGGV
R. rubrum         ETGGGRRRLYLHNDEEAAP-----------FAQEPGTFTPWEDLP-ESDLLFYEGLHGAV
R. sphaeroides    ETGQGRTRTYVHDDAEAAR-----------TGVAPGNFTDWRDFDSDSHLLFYEGLHGAV
A. cryptum        ATGTGQYRHYIHDQDEAER-----------YGGTPGTFTPWEDLPAPTDILLYEGLHGAV
X. flavus         ATGSGRFRHYVHDAGEAKL-----------YNTEPGRFTDWEDLEQGTDILFYEGLHGAV
N. hamburgensis   ESGTGNTRYYVHDDVESAK-----------HGVPPGTFTDWQALPENSDLLFYEGLHGAV
N. vulgaris       ETGTGNTRYYVHDDAESAL-----------HGVPPGTFTDWQPLPDASDLLFYEGLHGAV
R. palustris DX1  ETGTGRTRNYVHDDEEAAL-----------HGVPPGNFTGWRDLDPQSDLLFYEGLHGAV
R. palustris Bis  ETGTGKTRYYVHDDKEAAI-----------HGVQPGNFTDWQTLPEGSDLLFYEGLHGAV
M. capsulatus     ETGTARRRFYVHNEAESQRL----------GGYKPGTFTPWEEVPAGTDLLFYEGLHGGV
A. ferrooxidans   ESGHGQMRYYVHDELEAEL-----------RGSAPGTFTPWQDIPLGTDLLFYEGLHGGV
C. M. oxyfera     EAGVGKKRYYLHNAKEAAFHCKRLADAGVTCNADSGEFTPWEDIESNTDILFYEGLHGLV
T. denitrificans  ETGGGKKRYYIHSDEEADQHNKRLN-----TSLNPGEFTPWEDVPPGSDVLFYEGLHGLV
A. vinosum        ETGCGKKRYYIHSDEEARQLNARLG-----TSLNPGEFTPWEDLPAGTDLLFYEGLHGLV
P. lunula         QGKAVYKPIYNHDT---------------------GFKDPPELIE-PNKVMVFEGLHPIY
L. polyedrum      QGKAVYKPIYNHDT---------------------GNKDPPELIE-PNKVMVFEGLHPIY
B. natans         QGRAIKKPIYNHDT---------------------GAIDPVETIH-PNHIIIVEGLHPML
E. gracilis       RGETIAKPIYNHVN---------------------GTLDTPEEIA-PASIMIIEGLHPLL
P. parvum         AGKSVSKPIYNHVT---------------------GELDPYEDVD-PTPIVIFEGLHPMY
G. theta          EGKKVMKPIYNHVN---------------------GTLDEAEEIT-PTPIIIFEGLHPFY
T. pseudonana     EGKTIMKPIYNHVN---------------------GTLDTPEEIE-PTPVIIIEGLHPFV
O. sinensis       NGESIEKPIYNHVN---------------------GTLDTPETIV-PTPIVIIEGLHPMY
P. tricornutum    DGKTVEKPIYNHVN---------------------GTLDTPETIE-PTPIIIFEGLHPMH
D. lutheri        AGNSVMKPIYNHIT---------------------GELDAPEEIV-PTDIVIFEGLHPML
O. tauri          EGKSVDKPIYNHVT---------------------GVFDPAEKIE-SPEVLILEGLHPFA
M. commoda        EGKAVDKPIYNHVT---------------------GVFDPAEKIE-SPSILILEGLHPFA
P. sitchensis     EGKSDMKPIYNHVT---------------------GILDPAEQIN-PPKILVIEGLHPMF
P. patens         EGKSVEKPIYNHVT---------------------GLLDAPETIH-PPKILVIEGLHPMF
S. oleracea       EGKAVDKPIYNHVS---------------------GLLDPPELIQ-PPKILVIEGLHPMY
A. thaliana       NGIAVEKPIYNHVT---------------------GLLDPPELIQ-PPKILVIEGLHPMF
S. moellendorf.   EGKAVQKPIYNHVS---------------------GLLDPPELIQ-PPKILVIEGLHPMF
T. aestivum       EGKAIEKPIYNHVT---------------------GLLDPAELIQ-PPKIFVIEGLHPMY
P. trichocarpa    DGTAVEKPIYNHVT---------------------GLLDPPELIK-PPKILVIEGLHPMY
M. crystallinum   EGKAVEKPIYNHVT---------------------GLLDAPELIK-PPKILVIEGLHPMF
P. sativum        DGKSVQKPIYNHVT---------------------GLLDAPELIK-PPKILVIEGLHPMY
C. variabilis     EGKAVDKPIYNHVT---------------------GLLDPPEPTS-SPNILVIEGLHPFY
C. reinhardtii    EGKSVDKPIYNHVS---------------------GLIDAPEKIE-SPPILVIEGLHPFY
V. carteri        EGKAVDKPIYNHVT---------------------GLIDAPEKID-SPNILVIEGLHPFF
G. sulphuraria    EGYDIMKPIYNHQT---------------------GLIDPPELIQ-PNHIIVIEGLHPWY
G. kilaueensis    SGQSIQKPIYNHET---------------------GKIDPPERID-PTPIVILEGLHPFY
G. violaceus      VGQSIMKPIYNHET---------------------GKIDPPELIE-PTPIVILEGLHPFY
T. elongatus      NGESIMKPIYNHET---------------------GTIDPPEKVD-PNHVIVIEGLHPLY
Synechocystis sp  SGQSIMKPIYNHET---------------------GLLDPPEKVE-PNKVVVIEGLHPLY
M. aeruginosa     GGQAINKPIYNHET---------------------GMIDPPEIIE-PNKVIVIEGLHPLY
Cyanothece sp     EGQPIMKPIYNHET---------------------GMIDPPERIE-PNKVVVIEGLHPLY
S. elongatus      NGETIMKPIYNHET---------------------GLIDPPEKIE-PNRIIVIEGLHPLY
A. variabilis     EGQTINKPIYNHET---------------------GLIDPPEIVK-PNHIVVVEGLHPLY
N. spumigena      EGQVIQKPIYNHET---------------------GMIDPPERVE-PNHIIVVEGLHPLY
A. boonei         KGNAIIKPVYNHKT---------------------GKFDPPEVFE-PKKIVIVEGLHTLY
F. placidus       EWKEFEKPVYDHEK---------------------GEVKCCEIFK-PEKIVIAEGLHVLY
A. profundus      KGEKIKKPTYNHKT---------------------GTFGEWEDFE-STPIVIVEGLHTLY
A. veneficus      KGETIRKPTYDHST---------------------GTFGEWEDFT-PTPVVIVEGLHTLY
M. harundinacea   LGETIAKPVYDHSV---------------------GEFREPVPFR-SGPVLILEGLHPLY
M. concilii       RNERIDKPVYDHRT---------------------GEISGTVPFG-PAPVIIVEGLHPLY
M. thermophila    RGLAIEKPVYDHTT---------------------GEIRGPVIFK-PSPVIIVEGLHPFY
M. hungatei       AGRTIQKPVYLHDH---------------------GTFGEPELFS-PTKFIIIEGLHPYA
Methanolinea sp   QGKGIYKNIYNHST---------------------GRLEGPEYLA-PAKIIILEGLHSLF
M. boonei         QGYAIEKPVYNHAT---------------------GTFDPPIPFP-SKKILILEGLHTLF
M. limicola       SGKEILKPVYNHKT---------------------GKFDEPVPFS-SSKILIMEGLHAFA
M. petrolearia    EGKSIDKPVYNHDS---------------------GKIEGPVRLS-PSRIIIIEGLHPLF
M. palustris      EGRTIDKPVYNHAN---------------------GRFAPPIRFT-PGKILILEGLHTFF
M. marisnigri     AGRTIEKPVYNHDY---------------------GRFDPPVPFS-PTKILILEGLHPFI
M. liminatans     SGNTVMKPVYNHSD---------------------GTFDPPIPFR-PARVLILEGLHTLF
Gblocks
Annotation                   #  #                                  ########
                                                                  Walker B
```

646

647

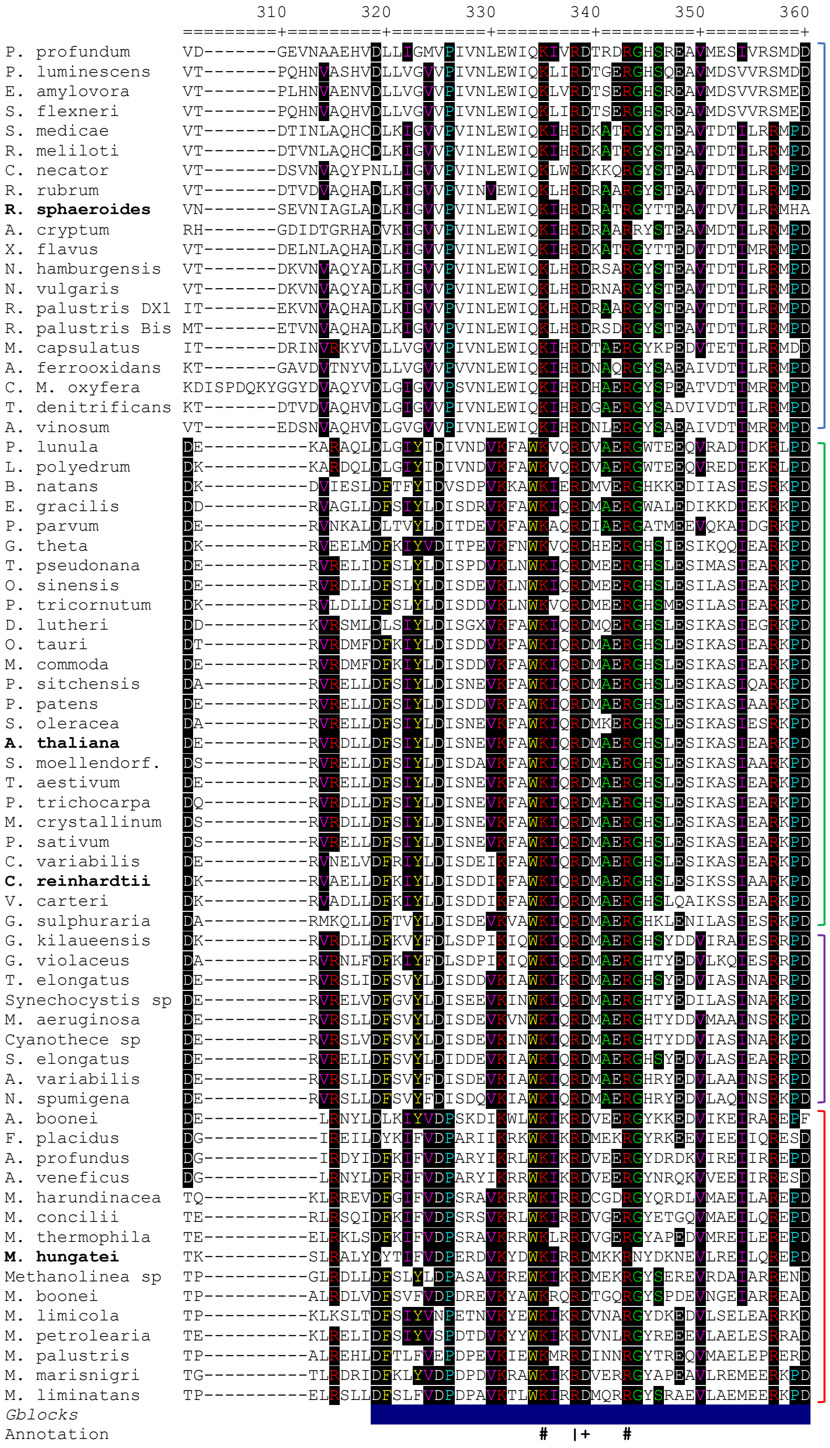
310
320
330
340
350
360
P. profundum
P. luminescens
E. amylovora
S. flexneri
S. medicae
R. meliloti
C. necator
R. rubrum
R. sphaeroides
A. cryptum
X. flavus
N. hamburgensis
N. vulgaris
R. palustris DX1
R. palustris Bis
M. capsulatus
A. ferrooxidans
C. M. oxyfera
T. denitrificans
A. vinosum
P. lunula
L. polyedrum
B. natans
E. gracilis
P. parvum
G. theta
T. pseudonana
O. sinensis
P. tricornutum
D. lutheri
O. tauri
M. commoda
P. sitchensis
P. patens
S. oleracea
A. thaliana
S. moellendorf.
T. aestivum
P. trichocarpa
M. crystallinum
P. sativum
C. variabilis
C. reinhardtii
V. carteri
G. sulphuraria
G. kilaueensis
G. violaceus
T. elongatus
Synechocystis sp
M. aeruginosa
Cyanothece sp
S. elongatus
A. variabilis
N. spumigena
A. boonei
F. placidus
A. profundus
A. veneficus
M. harundinacea
M. concilii
M. thermophila
M. hungatei
Methanolinea sp
M. boonei
M. limicola
M. petrolearia
M. palustris
M. marisnigri
M. liminatans
Gblocks
Annotation

```
                       370       380       390       400       410       420
                  =========+=========+=========+=========+=========+=========+
P. profundum      YLNYITPQFSRTHINFQRVPT-----------VDTS----NPFSAKGIPSLD--ESFVVI
P. luminescens    YINYITPQFSRTHINFQRVPT-----------VDTS----NPFSAKAIPSLD--ESFIVI
E. amylovora      YINFITPQFSRTHINFQRVPT-----------VDTS----NPFAARGIPSLD--ESFVVV
S. flexneri       YINYITPQFSRTHLNFQRVPT-----------VDTS----NPFAAKGIPSLD--ESFVVI
S. medicae        YVHYICPQFSLTDINFQRVPI-----------VDTS----NPFIARWIPTPA--ESILVI
R. meliloti       YVNYICPQFSLTDINFQRVPI-----------VDTS----NPFIARWIPTPA--ESILVI
C. necator        YVNYICPQFSRTHVNFQRVPC-----------VDTS----NPFISREIPAPD--ESMVVI
R. rubrum         YVHYICPQFTRTDVNFQRVPL-----------VDTS----NPFVARHVPSAD--ESFVVI
R. sphaeroides    YVHCIVPQFSQTDINFQRVPV-----------VDTS----NPFIARWIPTAD--ESVVVI
A. cryptum        YVHYICPQFTETDINFQRVPT-----------VDTS----NPFIARWIPTAD--ESIVVI
X. flavus         YVRYICPQFTETDINFQRVPT-----------VDTS----NPFVARWIPTPD--ESMVVI
N. hamburgensis   YVNYICPQFAETDINFQRVPT-----------VDTS----NPFIARWIPTPD--ESMVVI
N. vulgaris       YVNYICPQFAETDINFQRVPT-----------VDTS----NPFISRWIPTPD--ESMVVI
R. palustris DX1  YVHYIVPQFAETDINFQRVPT-----------VDTS----NPFIARWIPTAD--ESMVVI
R. palustris Bis  YVHYIVPQFGETDINFQRVPT-----------VDTS----NPFIARWIPTPD--ESMVVI
M. capsulatus     YVKVITPQFSQTDINFQRVPT-----------VDTS----NPFIARDIPTPD--ESFVII
A. ferrooxidans   YIHYITPQFSRAHINFQRVPL-----------VDTS----NPFIARDIPTPD--ESMVVI
C. M. oxyfera     YINHLTPQFSRTHVNFQRVPT-----------VDTS----NPFIARDIPSAD--ESFVVI
T. denitrificans  YVNYITPQFSRTDINFQRVST-----------VDTS----NPFISRDIPTPD--ESFIVI
A. vinosum        YIRYITPQFSLTDINFQRVPT-----------VDTS----NPFIARDIPTPD--ESFVII
P. lunula         FSAYVDPQKANADVILRYEPS-----------DQGL----PYLKVKLIQKKG--GKFPPI
L. polyedrum      FSAYVDPQKADADVILRYEPS-----------DQGL----PYLKVKLIQKKG--GAFPPI
B. natans         FEKFVEPQKANADVIISIEPTKLDVA------PGEET---KYLNTRLIQKENQ-HGIRPV
E. gracilis       FDKYVAPQRAKADMVIEVLPSRLAPP------KDETAPL-EYLRVRLIQKTTT-KHFDPV
P. parvum         FAAYVEPQKAKADIIVQVLFSDL-I-------DDPTG---KFLKVRLITKNNL-KHISPA
G. theta          FDAYIDPQKNKADCVIQVLPTNL-VA------NDK-----THLNVKLIQCKGV-DHYAPT
T. pseudonana     FDAFIEPQKKFADYVIEVLPTDL---------DKEDK---KTLKVRAIQKKGV-ADFTPT
O. sinensis       FDAYIAPQKEFADLTIEVLPTQL---------DEEDK---KTLRVRCIQKEGV-SDFSPC
P. tricornutum    FDAYIDPQKQLADLIIEVLPTRL---------DQDDK---KTLRVRCIQKEGV-ENFDPC
D. lutheri        FDAFVAPQRANADIVIEVLPTQL-V-------NDAEG---KFLRVRFIQKAGL-DLIKAP
O. tauri          FDAFVDPQKEHADVVIEVLPTQL-IP------DDNEG---KILRVRMIMKENV-ENFDAP
M. commoda        FDEFVDPQKQYADIVIQVLPTQL-IP------DDNEG---KILRVRMIMKEGV-ENFDAP
P. sitchensis     FDAFIDPQKQYADVVIQVLPTEL-IP------EENEG---KVLRVRMVMKEGV-NFFNPV
P. patens         FDAYIDPQKQYADVVIQVLPTQL-IP------DDNEG---KVLRVRMVMKEGV-PFFEPV
S. oleracea       FDAYIDPQKQHADVVIEVLPTEL-IP------DDDEG---KVLRVRMIQKEGV-KFFNPV
A. thaliana       FDAFIDPQKQYADAVIEVLPTTL-IP------DDNEG---KVLRVRLIMKEGV-KYFSPV
S. moellendorf.   FDAYIDTQKQYADVVIQVLPTQL-IP------DDNEG---KVLRVRMIMKEGV-DNFEPV
T. aestivum       FDAFIDPQKQYADAVIEVLPTQL-IP------DDNEG---KVLRVKLIMKEGI-KFFNPV
P. trichocarpa    FDAYIDPQKQYADAVIEVLPTQL-IP------DDNEG---KVLRVKLIMKEGV-EFFSPV
M. crystallinum   FDAYIDPQKQYADAVIEVLPTQL-IP------GDNEG---KVLRVRLIQKEGV-QYFSPV
P. sativum        FEAYIDPQKQYADAVIEVLPTQL-IP------DDNEG---KILRVRLIQKAGV-KYFSPV
C. variabilis     FDAYIDPQKKKADMIIQVLPTQL-VP------DEKEG---KILRVRLIMKDGK-KLFDPV
C. reinhardtii    FDAYIDPQKKDADMIIQVLPTQL-VP------DDK-G---QYLRVRLIMKEGS-KMFDPV
V. carteri        FDAYIDPQKKDADMIIQVLPTQL-VP------DDK-G---QYLRVRLIQKEGS-KMFDPV
G. sulphuraria    FQQYIDPQKKDAVAVIQVLPTRL-IP------DDTEK---KVLRVRLIQREGI-QGFQSV
G. kilaueensis    FSAYIDPQKQHADVVLQILPSELPEK------KDGT----KKIKAAMVQVEGI-PNYDPP
G. violaceus      FSAYIDPQKQYADVVLQILPSEL---------PEKAGGI-KRVKACMVQVDGI-PNYDAP
T. elongatus      FMAYIDPQKQYADVVLQILPSQL-AK------EEKVG---NILRVRMLQREGI-PGFEPV
Synechocystis sp  FTAYIEPQKQYADVVIQVLPTRLI--------EDKES---KLLRVRLVQKEGV-KFFEPA
M. aeruginosa     FSAYIDPQRQYADVVIQVLPTKLL--------EDHES---KLLRVRLIQKEGV-ENFEPA
Cyanothece sp     FSAYIEPQREYADVVIQVLPTNL-IE------NDKES---KILRVRLIQKEGI-ENFQPV
S. elongatus      FKAYIEPQRGHADIVIRVMPTQL-IP------NDTER---KVLRVQLIQREGR-DGFEPA
A. variabilis     FQKYIEPQREFADVVLQVLPTNL-IK------DDTDR---KVLRVRMLQREGK-EGFEPA
N. spumigena      FEKFIEPQREFADVVLQVLPTNL-IK------NDTER---KVLRVRMLQREDK-EGFEPT
A. boonei         YKRYIDFQKIYADIVIKIDISKFS--------EEDSYSVETILKVLDFPLSGIEMPLSLS
F. placidus       YKRYVDFQKIYADVVIKIYPSSLESTERISTLLGME----EMYRVRLIFTNLS-FELDHV
A. profundus      YKRYIDFQKIYADVVIKILPSSIQSTERIKYLVDSRE---EVYKVRLILKKTD-IPLESV
A. veneficus      YKRYIDFQKIYADVVIKIFPTALQSVERITYLTEAPP---ELYKVRLILKSGDFVPTEPI
M. harundinacea   YKLYVDVQKVYAEAVVKIDDTRLGAPSF----LEDPG---ERYSVRLIQEVLE-AGPSAV
M. concilii       YKLYVDIQKIYAQIVIKIQDSRF-HPSL--LDSQSTL---DWYSVRLIMEIME-HPVSEV
M. thermophila    YKLYVDVQKVYAEMVIKIQDSRI-PPEWTEIARGSAR---EKYSVRIIQQILD-HPLDEV
M. hungatei       YFQYVFPQREVADAVIQISYSSYGKE------EGEKR---NVYRVMLSMPAQE-YCFEDI
Methanolinea sp   YRAYIAPQRDHADAVIEIGFSRFGRD------LGWKK---NIYRVSILQTATS-CSMDHI
M. boonei         YATYIEPQRCNADAIIRIGYSKYGKD------LGPAK---NVYSISLLQNPMD-QAVRNT
M. limicola       YENFVLPQSESADALIEISDSSFDSP------SLKDR---GVYKITLYQKRLD-KTVKNI
M. petrolearia    YENYVHPQIRFADAVIGISKSRY-AD------IMNER---GVYRVVLYQKKQD-RTIRNI
M. palustris      YQRFIAPQQKYADVVVRVRFSKYGRE------RGIRE---KIYQVSLSQNRIT-KSIEDV
M. marisnigri     YERYVAPQCLFADAVIRIAFSKYGRD------VSEKR---NVYRVTLCQSRLD-KSIGDV
M. liminatans     YERYIAPQRAYADAVIRIACSGY---------GEEASEERNVYRVTILQKKLL-QQMVEI
Gblocks
Annotation
```

```
                          430       440       450       460       470       480
                 =========+=========+=========+=========+=========+=========+
P. profundum     RFRG-------------IEN---VDFP---YLLSMIQGSFMSRHNTLVVPGGKMS-----
P. luminescens   RFRD-------------LTQ---IDFP---YLLAMLQGSFVSSINTIVVPGGKMG-----
E. amylovora     HFQG-------------LED---IDFP---YLLSMLQGSFISHIKTLVVPGGKMG-----
S. flexneri      HFRN-------------LEG---IDFP---WLLAMLQGSFISHINTLVVPGGKMG-----
S. medicae       RFAK-------------PQS---IDFP---YLLSMLHNSYMSRANSIVVPGDKLD-----
R. meliloti      RFAK-------------PQS---IDFP---YLLSMLHNSYMSRANSIVVPGDKLD-----
C. necator       RFAN-------------PKG---IDFQ---YLLSMIHDSFMSRANTIVVPGGKME-----
R. rubrum        RFRD-------------PKG---IDFP---YLLNMLNDSFMSRPNTIVVPGGKME-----
R. sphaeroides   RFRN-------------PRG---IDFP---YLTSMIHGSWMSRANSIVVPGNKLD-----
A. cryptum       RFRD-------------PHG---IDFP---YLLAMLHGSFMSRANSIAVPGNKFD-----
X. flavus        RFRD-------------PHG---IDFP---YLLSMIHNSFMSRANSIVIPGNKQD-----
N. hamburgensis  RLKN-------------PRG---IDFP---YLLSMIPNSFMSRANSIVIHGSKMD-----
N. vulgaris      RLKN-------------PRG---IDFP---YLLSMIPSSFMSRANSIVIHGSKLD-----
R. palustris DX1 RFKN-------------PRG---IDFA---YLLSMIQGSFMSRANSIVIHGAKLD-----
R. palustris Bis RFKS-------------PRG---IDFA---YLLSMIQGSFMSRANSIVIHGSKMD-----
M. capsulatus    RFKE-------------PGKFN-VDFP---YLLAMLQNSFMSRHNSIVIPGGKMG-----
A. ferrooxidans  RFRD-------------PKE---ENFP---TLLQMLPGSFMSRSNTLVIPGTKMG-----
C. M. oxyfera    RFVD-------------PKRFG-VDFP---TLLVMINGSFISRRNTIVVPGGKMV-----
T. denitrificans RFRN-------------PQG---VDFP---YLLNMICNSFMSRRNTIVVPGGKMG-----
A. vinosum       RFKD-------------PKKLQ-IDFP---YLLSMIHDSFMSRRNSMVVPGGKMG-----
P. lunula        SLKK-------------DLSLT-GSKP--GATMKMYDDDWFGNPVTVVEMDGEIDMDNMA
L. polyedrum     SLKK-------------DLTLT-GSKP--GATLKMYDDDWFGNAVTVVEMDGEIDMDNME
B. natans        YMFEEEGSTVDWDPCAGPGAMA-CPYP--GTRVRYYNEMSGEKHAHVLEVDGVFG---ET
E. gracilis      YLIEKG-SSVTWKPC--GDNLQ-CEYP--GLQLAYYTEEYMGHPAEVLEMDGVIH---NL
P. parvum        YLMDEG-ASITWKPN--PNKLT-TSAP--GVLFKSYQDEWFGQSVSVLEMDGKID---SL
G. theta         YLWDEG-SDIEWVPP--RNKLA-SSAPGAGLKIYQKTEKWAGKDAAVIGMDGKYD---KI
T. pseudonana    YLFDEG-SEIEWAPS--ADKLS-SPAP--GIKLSYKQEQYFGADVAVVEMDGTFD---NI
O. sinensis      YLFDEG-STIAWTPA--PSKLS-SSGP--GLTMAYGTEDYYGKPAQVVEMDGTFD---NI
P. tricornutum   FLFDEG-SSIEWTPA--PTKLS-SPAP--GIKLAYYPEEFFGKDAQVLEMDGNFD---NI
D. lutheri       FLFDEG-STIEWTPC--GKKLT-CAYP--GIKFRYGTEMYMGSEVTVLEMDGRFD---KL
O. tauri         YLFDEG-STISWIPC--GRKLT-CSYP--GIKFFYGPDTYYGKEVTVLEMDGQFD---KL
M. commoda       YLFDEG-STISWIPC--GRKLT-CSYP--GIKFFYGPDTFFGEEVSVLEMDGQFD---KL
P. sitchensis    YLFDEG-STISWIPC--GRKLT-CSYP--GIKFFYGPDIYYDNEVSVLEMDGQFD---RL
P. patens        YLFDEG-STISWIPC--GRKLT-CSYP--GIKFFYGPDTYYGNEVSVLEMDGQFD---KL
S. oleracea      YLFDEG-STISWIPC--GRKLT-CSYP--GIKFSYGPDTFYGNEVTVVEMDGMFD---RL
A. thaliana      YLFDEG-STISWIPC--GRKLT-CSYP--GIKFNYEPDSYFDHEVSVLEMDGQFD---RL
S. moellendorf.  YLFDEG-STISWIPC--GRKLT-CSYP--GIKFFYGPDTYYDNEVSVLEMDGQFD---KL
T. aestivum      YLFDEG-STINWIPC--GRKLT-CSYP--GIKFSYGPDTYFGQEVSVLEMDGQFD---RL
P. trichocarpa   YLFDEG-SSISWIPC--GRKLT-CSYP--GIKFSYGPDAYYGHEVSVLEMDGQFD---RL
M. crystallinum  YLFDEG-SSITWIPC--GRKLT-CSYP--GIKFFYGPDTYFGNEVTVLEMDGQFD---RL
P. sativum       YLFDEG-STISWIPC--GRKLT-CSYP--GIKFFYGPETYKGNEVSVVEMDGQFD---RL
C. variabilis    YLFDQG-STVSWIPC--GRKLT-CSFP--GIKMFYGPDTYYGEEVSVLEMDGQFD---KL
C. reinhardtii   YLFDEG-STISWIPC--GRKLT-CSFP--GIKMFYGPDTWYGQEVSVLEMDGQFD---KL
V. carteri       YLFDEG-STISWIPC--GRKLT-CSFP--GIKMFYGPDTWYGQEVSVLEMDGQFD---KL
G. sulphuraria   YLYDEG-STIDWIPC--GRKLT-CSYP--GIKFHYGPDNWYNHDVSVLEVDGNFE---KL
G. kilaueensis   YLFDRAISDVCW-----KPNFP-NSDP--ELTFCYGTTDYFGRPASYLSIDGEFA----P
G. violaceus     YLFDRAISDVCWK----PNFPN-SDEE---LTFCYGTQDYFGRPASYLSIDGEFA----P
T. elongatus     YLFDEG-STITWIPC--GRKLT-CSYP--GIRLSYGPDEYYGHPVSVLEVDGRFE---KL
Synechocystis sp YLFDEG-STIDWRPC--GRKLT-CTYP--GIKMYYGPDNFMGNEVSLLEVDGRFE---NL
M. aeruginosa    YLFDEG-STIDWRPC--GRKLT-CTYP--GIKLYYGPDGFLGNEVSILELDGQFD---NL
Cyanothece sp    YLFDEG-STIDWRPC--GRKLT-CAYP--GLKMYYGPDNIFGNEVSILEIDGQFD---NL
S. elongatus     YLFDEG-STIQWTPC--GRKLT-CSYP--GIRLAYGPDTYYGHEVSVLEVDGQFE---NL
A. variabilis    YLFDEG-STINWTPC--GRKLT-CSYP--GMQLYYGSDVYYGRYVSVLEVDGQFD---NL
N. spumigena     YLFDEG-STINWTPC--GRKLT-CSYP--GMQVYYGSDVYYGRYVSVLEVDGQFD---NL
A. boonei        SLINT------------------SQKP---MMISYRDDFYYMKRVSRITFDGLIP-----
F. placidus      TLNVDL-----------SRLIK-ASER--DFAISFFSDYYYGKRASFIDIDGFLN-----
A. profundus     SLNIDI-----------SDLVK-ASER--DFSIGFFSDYYYAEKASFIDIDGFLN-----
A. veneficus     KLDIDL-----------SAFVR-ASEK--DFALGFFSDYYYEKPASFIDIDGLMN-----
M. harundinacea  DLKIDL-----------SAL---FSLPDGEFSISFQRDDYYGKRVSVLTVDGELH----R
M. concilii      SMNIDL-----------SKILR-RSEH--EFSIEFQRDDYYGKRVGIMTMDGEIH-----
M. thermophila   DLTIDL-----------SRIMR-LTER--EFSIEFQRDDYYGKRVGVMTLDGEFP----L
M. hungatei      ELNIDL-----------CDLFK-KSSH--DFSLSCISHTPDSRNMRALVVDGELM-----
Methanolinea sp  CLGMDL-----------VSLMS-SFDH--NFMVEYRVVTEAGRRMGRLAFDGGIP-----
M. boonei        SLSIDL-----------TSLLA-FHES--EFSIEFSTQRIECSFLRSLTFDGEMN-----
M. limicola      CLNFDL-----------FAINS-LADR--NFRFDFRVTERGGEKIGALSLDGEFQ----Y
M. petrolearia   NLNFDL-----------FAINS-LAER--GFSFDFSIIEKYEKKMGALSLDGEFR-----
M. palustris     DLSIDL-----------FSLLS-LSDR--NFLIEFSHEQRNDERTGELILDGELS-----
M. marisnigri    DLSIDL-----------FGLLS-LSER--DFMVEFTIEDVGGEAMGALTFDGELN-----
M. liminatans    GLSIDL-----------GEIFRLCERP---FLLEFGLSSLDGRELSALVLDGELA-----
Gblocks
Annotation                          *        *
                                 C243     C249
```

```
                        490       500       510       520       530       540
                 =========+=========+=========+=========+=========+=========+
P. profundum     -----------------------------------FAMELIVRPLIQQLIDTGKVG----
P. luminescens   -----------------------------------LAMELIMTPLVQRLLEGGKIS----
E. amylovora     -----------------------------------LAMELIMGPLVKRLIEGKKIG----
S. flexneri      -----------------------------------LAMELIMLPLVQRLMEGKKIE----
S. medicae       -----------------------------------LAMQLIFTPLIHKLLERKHRMS---
R. meliloti      -----------------------------------LAMQLIFTPLIHKLLERKHRMS---
C. necator       -----------------------------------LAMQLIFTPFVLRMMERRKRAAQ--
R. rubrum        -----------------------------------LSMQLIFTPFIWRFMDKRARALGR-
R. sphaeroides   -----------------------------------LAMQLILTPLIDRVVRESKVA----
A. cryptum       -----------------------------------LAMQLILTPIIMQLMDRKRRAQ---
X. flavus        -----------------------------------LAMQLLLTPLIMKLMDRKRRAG---
N. hamburgensis  -----------------------------------LAMQLILTPLILQLIDRKRRA----
N. vulgaris      -----------------------------------LAMQLILTPLILQLIERKKRA----
R. palustris DX1 -----------------------------------LAMQLILTPLIMQLIDRKRSVK---
R. palustris Bis -----------------------------------LAMQLILTPLIMQLIERKRGAK---
M. capsulatus    -----------------------------------LAMEIIFRPILERMMADREKG----
A. ferrooxidans  -----------------------------------YAMELILGPRIERMLEDMHLTI---
C. M. oxyfera    -----------------------------------LAMELILAPIVHIMIENKLKA----
T. denitrificans -----------------------------------FAMELILAPLIQDMIANKKK-----
A. vinosum       -----------------------------------FAMEIILQPIIERMMDARRV-----
P. lunula        DQLKEIEENLEGLAGS-PGELTEAMTKLRSSPGSQNGTGLLQTVIAMKVREVYERLTAKV
L. polyedrum     AQLKEIEDNLEGLPSKTPGELTEAMVKLKSSPGSQNGTGMLQTVIAMKFREVYEKLTGGK
B. natans        EELFFIEERLSNTNTKYFGELTKQMLKNKAAPGSGDGSGLIQALVALKMRESYERFTGRT
E. gracilis      KEGLYVEKFLHNTGAKEFGELTQELLKGQNSPGGDNGTGFMQTLAALKIREIYERATGE-
P. parvum        EELIYVESQLCNTGTKYYGELTEQMVNNKASPGSENGSGLFQTICAFKIREAFEAITA--
G. theta         DEMMYVEKQFASTGSKFVGEITKKMLEYEGQPGSNDGTGFLQTITALKVREVYEGIAKVK
T. pseudonana    QELVYVESNLGNTNSKFYGEVTQAMLSLADSPGSNNGTGLMQTLAAFAIRELYNKKSAAA
O. sinensis      QELVYVESQLSNTSTKFYGELTQAMLKLADAPGSNNGTGLMQTLAAFAIRELYEKKAAAA
P. tricornutum   QELVYVESALSNTKTKFYGEMTQAMLALATAPGSNNGTGLMQTLAAFAIRDIYEKKTAAA
D. lutheri       DELIYVESALTNTGAKFYGELTQQILKNKDAVGSDNGTGFFQTLCSFKIREAYEKATGKT
O. tauri         EELIYVESHLSNTSSKFYGEITQQMLKYQNGPGSNNGTGFFQTIVGLKVREVYERISGKE
M. commoda       EELIYVESHLSNTSTKFYGEITQQMLKYQNGPGSNNGTGLFQTLCGLKVRELYERISEKE
P. sitchensis    EELIYVESHLSNISTKFYGEVTQQMLKHADFPGSNNGTGLFQTIVGLKIRDVYEQLISQT
P. patens        DELIYVESHLSNISTKFYGEITQQMLKHADFPGSNNGTGLFQTICGLKIRSVYERILANQ
S. oleracea      DELIYVESHLSNLSTKFYGEVTQQMLKHQNFPGSNNGTGFFQTIIGLKIRDLFEQLVASR
A. thaliana      DELIYVESHLSNLSTKFYGEVTQQMLKHADFPGSNNGTGLFQTIVGLKIRDLYEQLIANK
S. moellendorf.  DELIYVESHLSNISTRFYGEITQQMLKHADFPGSNNGTGLFQTIVGLKIRDVFERITSKQ
T. aestivum      DELIYVESHLSNLSTKFYGEVTQQMLKHADFPGSNNGTGLFQTIVGLKIRDLYEQIIAER
P. trichocarpa   DELIYVESHLSNISTKFYGEVTQQMLKHADFPGSNNGTGLFQTIVGLKIRDLFEQIVASR
M. crystallinum  DELIYVESHLSNLSTKFYGEVTQQMLKHQDFPGSNNGTGLFQTIVGLKIRDLFEQLIASK
P. sativum       DELIYVESHLSNLSSKFYGEVTQQMLKHADFPGSNNGTGLFQTIVGLKIRDLFEQIVAS-
C. variabilis    EELIYMESHLSNTSAKFYGEITQQMLKNSSFPGSNNGTGLFQTIVGLKCREVYERITQK-
C. reinhardtii   EELIYVESHLSNTSAKFYGEITQQMLKNSGFPGSNNGTGLFQTIVGLKVREVYERIVKKD
V. carteri       EELIYVESHLSNTSAKFYGEITQQMLKNSGFPGSNNGTGLFQTIVGLKVREVYERIVKKD
G. sulphuraria   EELIYIESHLNNTSTKFYGEITQQLLRNSSAPGSNNGTGLFQVLTALKMRQLYEELTGKS
G. kilaueensis   DDSPELEQKLRNTSETRFGELASLILKHTDYPGSRDGTGFFQVLIGLKFRSIYEALIAQS
G. violaceus     EDSPELEEKLRNTSETRYGELASLILKHTDYPGSRDGTGFFQVLMSLKFRAIYEALTAQL
T. elongatus     DELIYIESHLSNTSTKHYGEVTELLLKHRDYPGSDNGSGLFQVLTGLKMRATYERLTSRD
Synechocystis sp EEMVYVENHLSKTGTKYYGEMTELLLKHKDYPGTDNGTGLFQVLVGLKMRKVYEQLTAEA
M. aeruginosa    EEMIYIESHLSKTGTKYYGEMTELLLKHKDYPGSTNGTGLFQVLVGLKMRETYEKLMAAE
Cyanothece sp    EEMIYIEQHLSRTGTKYYGEMTHLLQQHKDYPGSNNGTGLFQVLVGLKMRETYEILTSV-
S. elongatus     EEMIYVEGHLSKTDTQYYGELTHLLLQHKDYPGSNNGTGLFQVLTGLKMRAAYERLTSQ-
A. variabilis    EEVIYIETHLSNTSTKYQGELTQLLLQHREYPGSNNGTGFFQVLTGLKMRAAYERLTAKE
N. spumigena     DEVIYVETHLSKTSTKYEGEMTHLLLQHREYPGSNNGTGLFQVLTGLKMRAAYERLTTK-
A. boonei        -------HEAIDSLERRIMEYTGFENYILERSKYVNGTQIAQLLVAWYFVEMMNNIFREI
F. placidus      ---VEIFESLLDALRDETGYKVRW-----EAREYVNSIEVAKLLLCWNLVEIIKMKG---
A. profundus     ---VDIFKSLFDSLRKEIGDG-----EIKVESEYVNAIEFSKLLVCWKLVEVLKHSLR--
A. veneficus     ---VELFSTLLDILKKEVGADVEWKMNT-----YVNAIEVAKLLVCWRFLELIKFRLEAE
M. harundinacea  SMVEGVEAKLKSLAG------STAPICDRAGGDRVTSTGMAQLLLSWRLLEKLQLHLDKA
M. concilii      ------QSMISDLEKKLGESLGTDGMISDRREEYVNAIGLAQLILTWNCVEKLDYLLQEE
M. thermophila   TMIKDLERKLCDFLGRDVPSVTEG--CQEDHGSYVNATEMTQLILIWRFLEKIANLAW--
M. hungatei      ------PDTIHKIERQIEFQTGISPINIFRGQEHITGTDLVRLILSWQIINGRIALSNHL
Methanolinea sp  ------REMAGDLRRKIEEQTGTQPADMFRGNAMVTATDMVRLIVSWRIINHLANLA---
M. boonei        ------YDTVRNLELSIEHQTGIHPIEMFAGRKVVTPTNIVQLLLSWRIINRRIFLQDHR
M. limicola      DVIRCLELNIEEQTG-------VSPVSLFEGRDYVTATEMIQLLLSWRIINRRIQMES--
M. petrolearia   ------HEVVSLLEKGIEMQTGIGPVSVYSDRSYVTATEMVELILSWRIINRRMDMKYGS
M. palustris     ------TRMVKRLETSIEEQTQVRPISDFHDHDYMTATEVVQLILAWRIIHQRVFLERCL
M. marisnigri    ------DAVARKLERNIEIQTQVEPIDLSQDSDYLTAGDMAQLILAWRIINRRIFIESAP
M. liminatans    ------ASAIRRLARTIGRETRSEAVNLFADRQYVTAGEIAELILAWRILNRWHVLEAAR
Gblocks
Annotation
```

```
                          550       560
                 =========+=========+
P. profundum     --------------------
P. luminescens   --------------------
E. amylovora     --------------------
S. flexneri      --------------------
S. medicae       --------------------
R. meliloti      --------------------
C. necator       --------------------
R. rubrum        --------------------
R. sphaeroides   --------------------
A. cryptum       --------------------
X. flavus        --------------------
N. hamburgensis  --------------------
N. vulgaris      --------------------
R. palustris DX1 --------------------
R. palustris Bis --------------------
M. capsulatus    --------------------
A. ferrooxidans  --------------------
C. M. oxyfera    --------------------
T. denitrificans --------------------
A. vinosum       --------------------
P. lunula        GA------------------
L. polyedrum     --------------------
B. natans        IQIPKWGLFLDDY-------
E. gracilis      KA------------------
P. parvum        --------------------
G. theta         VPAQAN--------------
T. pseudonana    KLAATKETAASA--------
O. sinensis      KLAVIEA-------------
P. tricornutum   KAKAGVSAAAA---------
D. lutheri       VDASAAA-------------
O. tauri         VVAKA---------------
M. commoda       VVTAA---------------
P. sitchensis    TGASLEAAKA----------
P. patens        KNAATLQSAKA---------
S. oleracea      STATATAAKA----------
A. thaliana      ATARAEAKA-----------
S. moellendorf.  GSPVGAAAATSKV-------
T. aestivum      AGVPAEAAKV----------
P. trichocarpa   AKTPVEATKA----------
M. crystallinum  TAAPAAATKA----------
P. sativum       RAETPVGAAKA---------
C. variabilis    --------------------
C. reinhardtii   VVPV----------------
V. carteri       VVPA----------------
G. sulphuraria   IAPVLV--------------
G. kilaueensis   LAAAKV--------------
G. violaceus     LTASKAK-------------
T. elongatus     AATVTNR-------------
Synechocystis sp KVPASV--------------
M. aeruginosa    AKVAASV-------------
Cyanothece sp    ESKVATQV------------
S. elongatus     AAPVAASV------------
A. variabilis    AKLAVQV-------------
N. spumigena     EAKLAVQV------------
A. boonei        ENAREKNIY-----------
F. placidus      --------------------
A. profundus     --------------------
A. veneficus     L-------------------
M. harundinacea  --------------------
M. concilii      GY------------------
M. thermophila   --------------------
M. hungatei      DQ------------------
Methanolinea sp  --------------------
M. boonei        --------------------
M. limicola      --------------------
M. petrolearia   GVSS----------------
M. palustris     HQDHNK--------------
M. marisnigri    GAGGTGRTVTGNNGHGCGRR
M. liminatans    SAHAKG--------------
Gblocks
Annotation
```

B

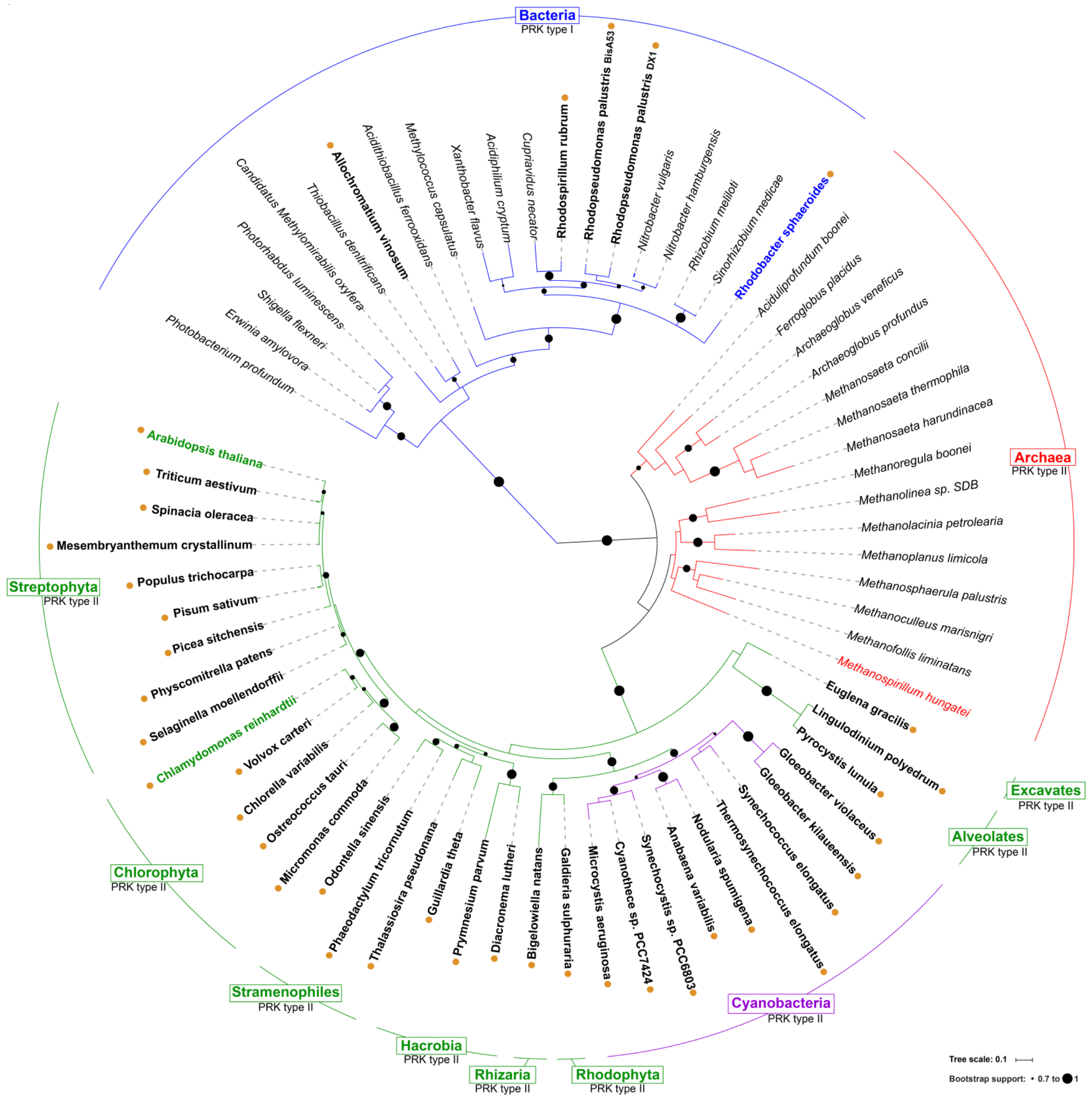

Bacteria
PRK type I
Rhodopseudomonas palustris BisA53
Rhodopseudomonas palustris DX1
Rhodospirillum rubrum
Cupriavidus necator
Acidiphilium cryptum
Xanthobacter flavus
Methylococcus capsulatus
Acidithiobacillus ferrooxidans
Allochromatium vinosum
Thiobacillus denitrificans
Candidatus Methylomirabilis oxyfera
Photorhabdus luminescens
Shigella flexneri
Erwinia amylovora
Photobacterium profundum
Nitrobacter vulgaris
Nitrobacter hamburgensis
Rhizobium meliloti
Sinorhizobium medicae
Rhodobacter sphaeroides
Aciduliprofundum boonei
Ferroglobus placidus
Archaeoglobus veneficus
Archaeoglobus profundus
Methanosaeta concilii
Methanosaeta thermophila
Methanosaeta harundinacea
Methanoregula boonei
Methanolinea sp. SDB
Methanolacinia petrolearia
Methanoplanus limicola
Methanosphaerula palustris
Methanoculleus marisnigri
Methanofollis liminatans
Methanospirillum hungatei
Archaea
PRK type II
Euglena gracilis
Lingulodinium polyedrum
Pyrocystis lunula
Gloeobacter violaceus
Gloeobacter kilaueensis
Synechococcus elongatus
Thermosynechococcus elongatus
Nodularia spumigena
Anabaena variabilis
Synechocystis sp. PCC6803
Cyanothece sp. PCC7424
Microcystis aeruginosa
Galdieria sulphuraria
Bigelowiella natans
Diacronema lutheri
Prymnesium parvum
Guillardia theta
Thalassiosira pseudonana
Phaeodactylum tricornutum
Odontella sinensis
Micromonas commoda
Ostreococcus tauri
Chlorella variabilis
Volvox carteri
Chlamydomonas reinhardtii
Selaginella moellendorffii
Physcomitrella patens
Picea sitchensis
Pisum sativum
Populus trichocarpa
Mesembryanthemum crystallinum
Spinacia oleracea
Triticum aestivum
Arabidopsis thaliana
Streptophyta
PRK type II
Chlorophyta
PRK type II
Stramenophiles
PRK type II
Hacrobia
PRK type II
Rhizaria
PRK type II
Rhodophyta
PRK type II
Cyanobacteria
PRK type II
Alveolates
PRK type II
Excavates
PRK type II
Tree scale: 0.1
Bootstrap support: 0.7 to 1

**Fig. S6. Sequence alignment and phylogenetic analysis of 69 PRKs.** (A) The sequences annotated as PRKs were retrieved from Uniprot and aligned using the phylogeny webserver suite (www.phylogeny.fr) (27). Blue areas in the "Gblocks" line define the conserved areas, later used by the software to determine the phylogeny. The alignment was performed by MUSCLE and curation by Gblocks. Annotations used are as follow: black boxes within the sequences indicate conservation of the residue in more than 70% of the sequences. Walker A (P-loop) and Walker B motives are represented by a red or green area, respectively, and the clamp loop is highlighted by a yellow area. Bars (|) or Hashtags (#) denote residues implicated in Ru5P or ATP binding, respectively. Plus sign (+) indicates two Aspartate residues shown to be crucial for catalysis by mutagenesis in *R. sphaeroides* (28). Star signs (*) indicate Cys residues implicated in disulfide bridges (Cys16 with Cys55; Cys243 with Cys249) (29), numbers below are for *C. reinhardtii* PRK. Blue and red bars on the right side indicate clusters of bacterial and archeal PRK, respectively, while the green one indicates the cluster of eukaryotic and the purple one is for the cyanobacterial PRKs. Species names in bold are indicating the 4 species with known structure. Uniprot accessions numbers and other details, for all protein sequences are reported in Table S6. (B) The phylogeny was built with PhyML and the tree with TreeDyn. The visual was obtained with iTOL (http://itol.embl.de/) (30). Bootstrap values superior to 0.7 are represented by black circles with a radius proportional to the value. Branch lengths are represented by straight lines at indicated scale, while dashes are presented for the sake of clarity. The clades are colored in function of their kingdom (Bacteria in blue, Archaea in red and Eukaryotes in green) except Cyanobacteria clade which is in purple. Photosynthetic species are in bold indicated by a yellow circle while the others are italicized. Species for which the PRK has a known structure, *i.e. R. sphaeroides* (31), *M. hungatei* (5), *C. reinhardtii* and *A. thaliana*, are represented in the color of their kingdom.

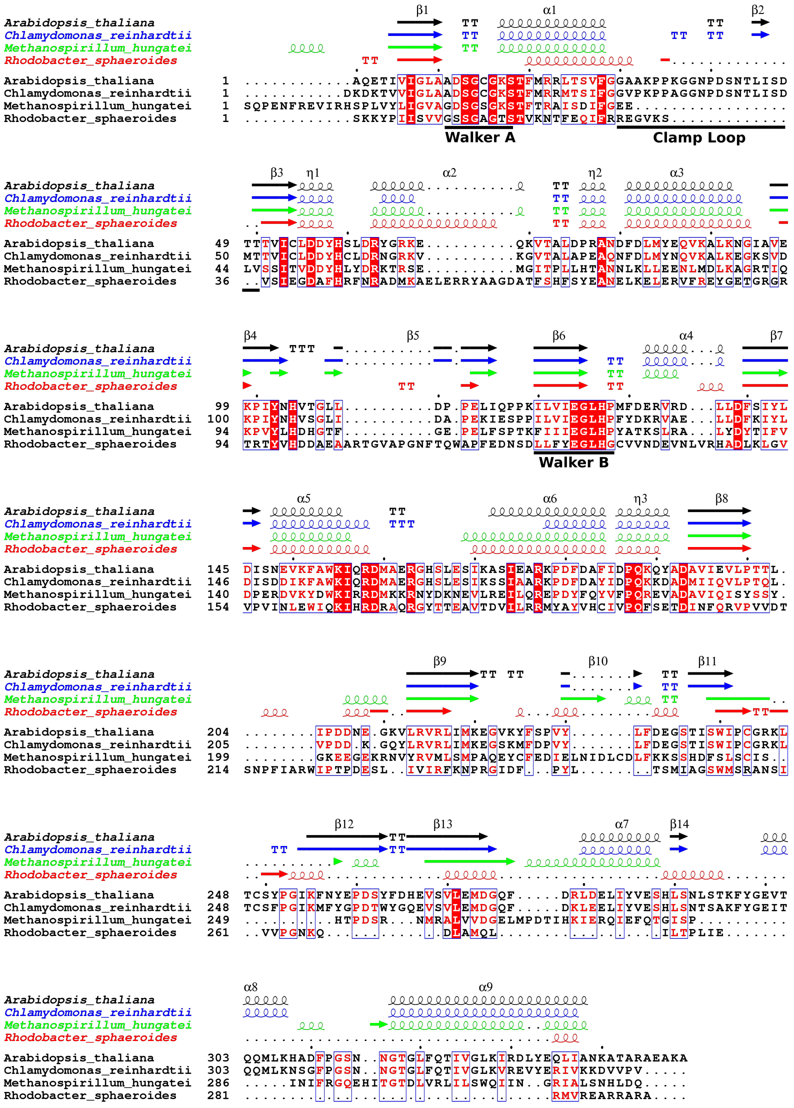


**Fig. S7. Sequence and structural alignment of the four structurally known PRKs.** The alignment was performed with Espript (http://espript.ibcp.fr) (32) using the sequence and the structure of *At*PRK (Uniprot accession code P25697 and PDB ID code 6H7H); *Cr*PRK (Uniprot accession code P19824 and PDB ID code 6H7G); *Rs*PRK (Uniprot accession code P12033 and PDB ID code 1A7J) (31), *Mh*PRK (Uniprot accession code Q2FUB5 and PDB ID code 5B3F) (5). The sequence of both photosynthetic PRKs is much longer (349 and 344 residues for *At*PRK and *Cr*PRK, respectively) than bacterial (290 residues) and Archae (323 residues) PRKs. Sequence identities among considered PRKs are: 75% for *At*PRK *vs* *Cr*PRK; 35% for *At*PRK *vs* *Mh*PRK; 32% for *Cr*PRK *vs* *Mh*PRK; 22% for *At*PRK *vs* *Rs*PRK; 24% for *Cr*PRK *vs* *Rs*PRK. The sequence identities were calculated by Clustal Omega (33).

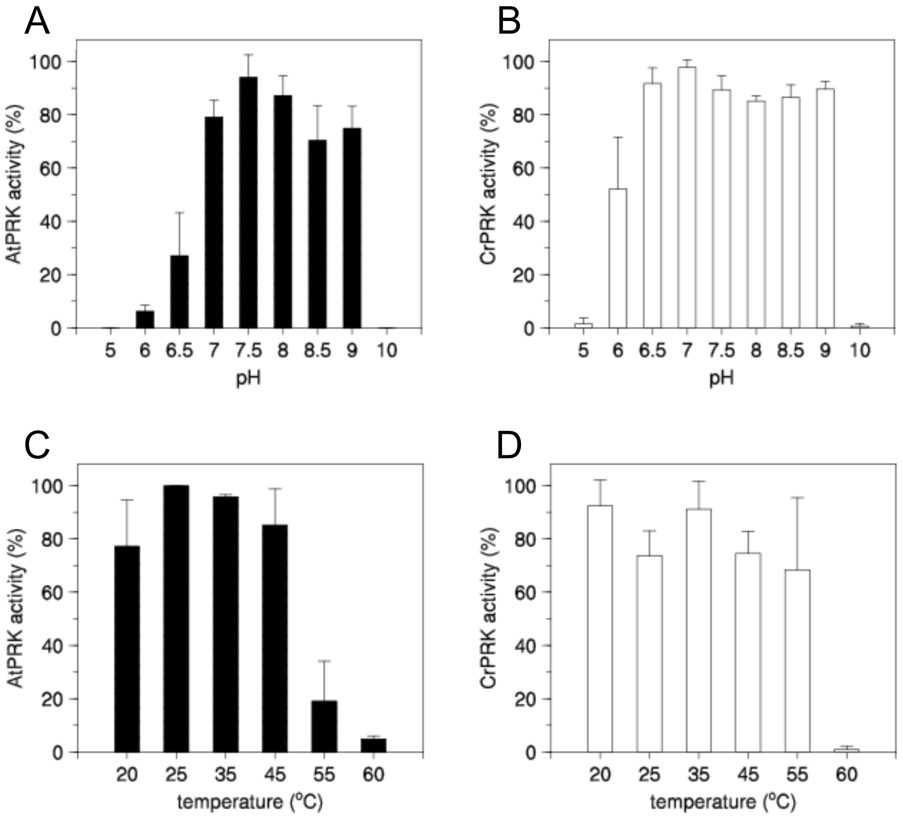


**Fig. S8. pH and temperature dependence of photosynthetic PRKs.** The enzyme activity of *At*PRK (A and C) and of *Cr*PRK (B and D) was evaluated at different pHs (upper panels) and temperatures (lower panels). Values are reported as means ± SD (n=3).

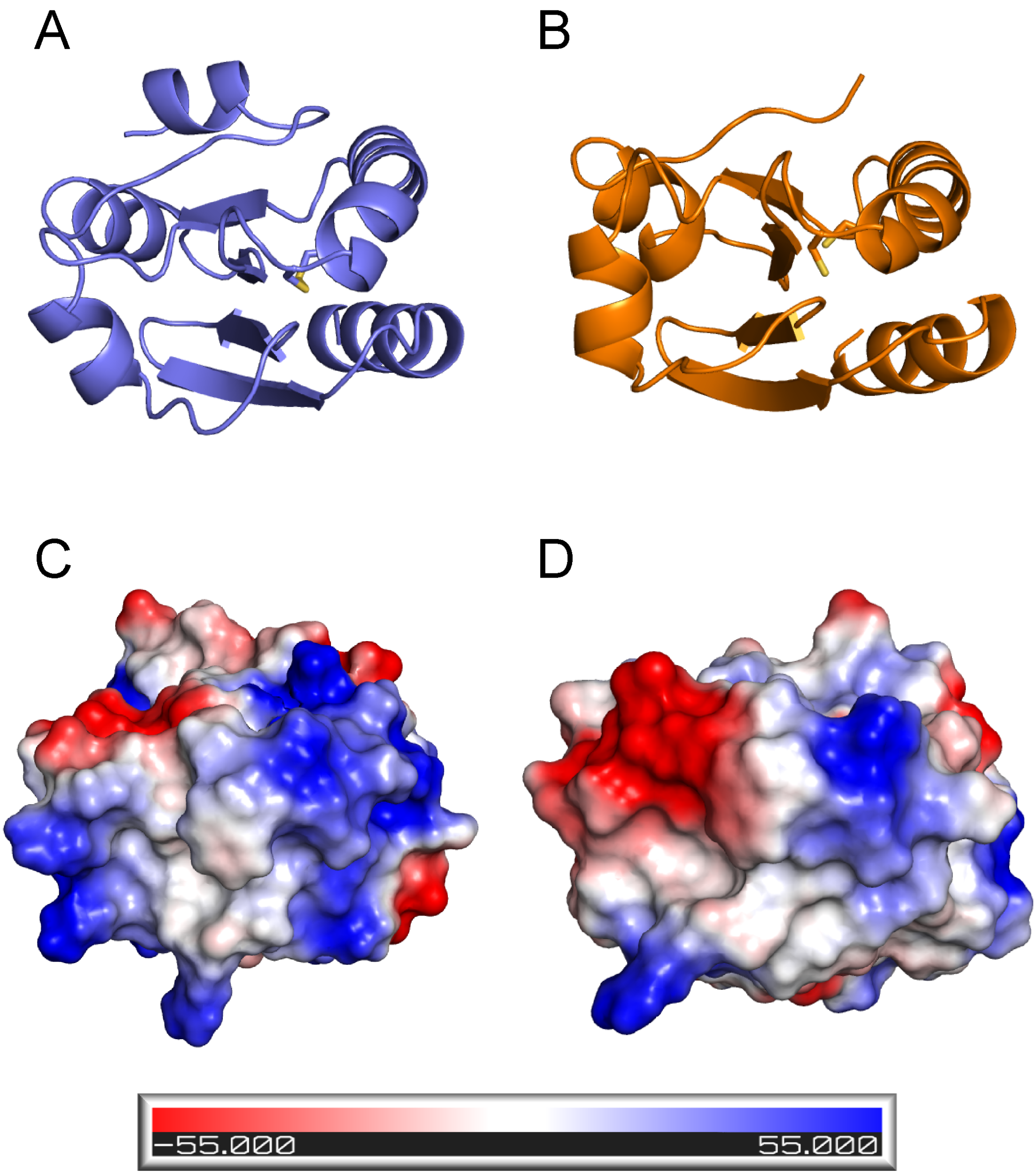


**Fig. S9. Electrostatic surface potential of *A. thaliana* TRX-f1 and TRX-m2.** Structure represented as ribbon (A, B) and electrostatic surface potential (C, D) of the homology model of *Arabidopsis thaliana* TRX-f1 (A, C) and TRX-m2 (B, D). The crystal structures of *Spinacia oleracea* TRX-f and TRX-m (PDB ID codes 1FAA and 1FB6) (34) were used as template to model *Arabidopsis thaliana* TRX-f1 and TRX-m2, respectively. The sequence identity of *Arabidopsis thaliana* TRXs with spinach enzymes is 59% and 75% for TRX-f1 and TRX-m2, respectively. The homology modelling was performed with Swiss-Model (35).

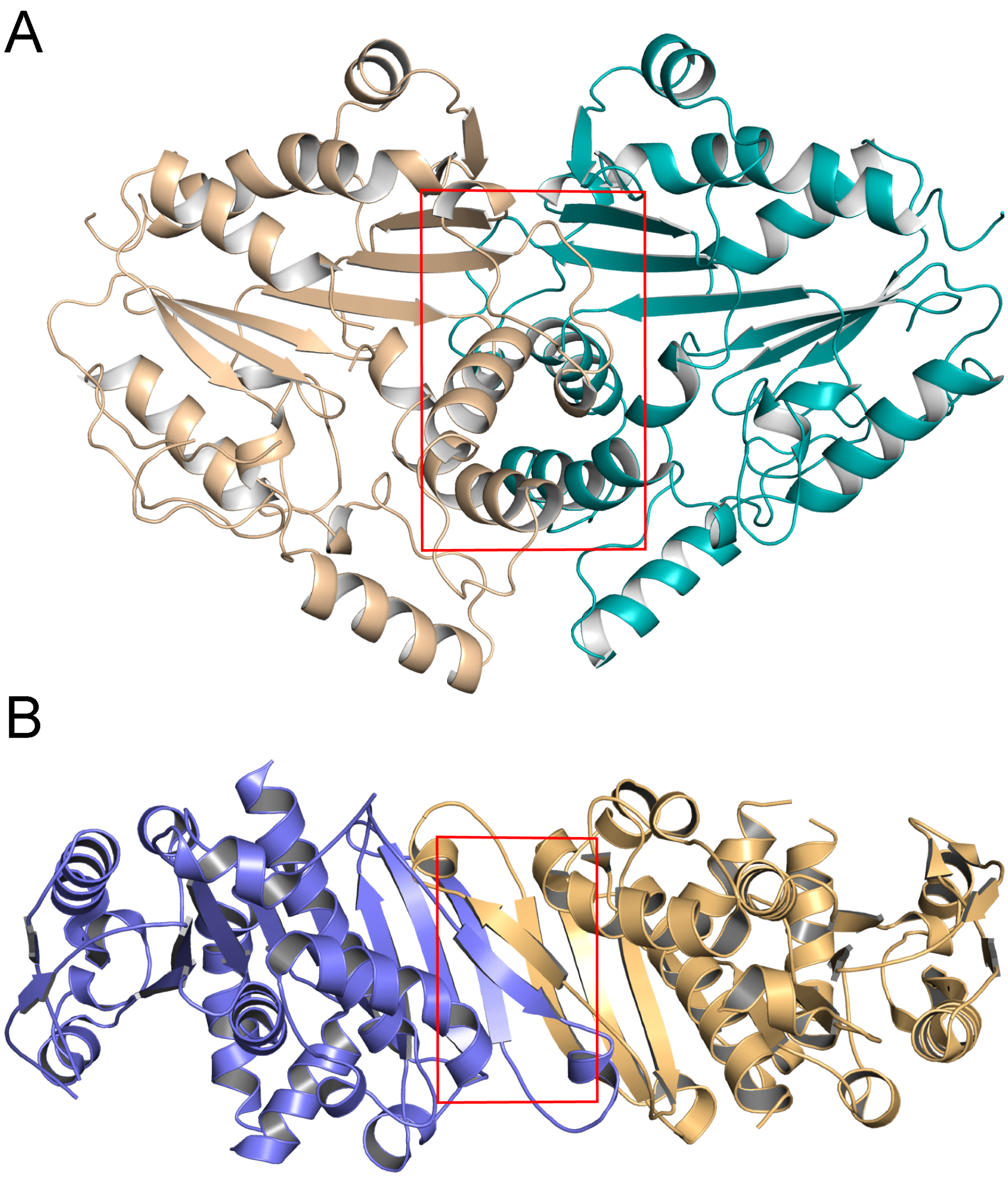


**Fig. S10. Dimer interface of bacterial and Archaea PRKs**. Dimer interface of (A), octameric *Rhodobacter sphaeroides* PRK (PDB ID code 1A7J) (31) and (B), dimeric *Methanospirillum hungatei* PRK (PDB ID code 5B3F) (5) is highlighted by a red box. The dimer interface of bacterial PRK is formed by three β-strands and one α-helix while in Archaea enzyme by two β-strands. The calculated dimer interface areas are 1667 and 1695 Å$^2$, respectively.

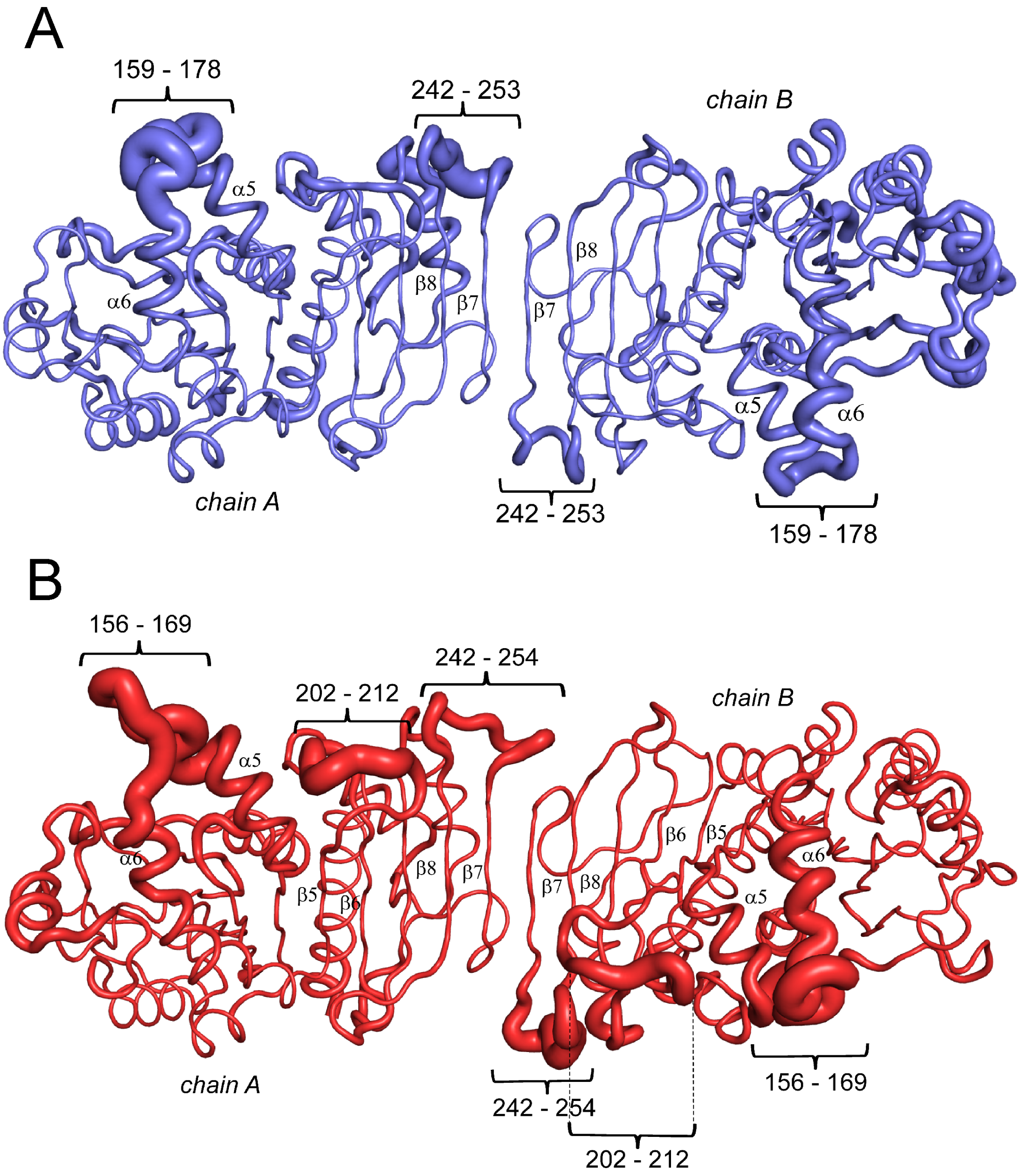


**Fig. S11. Flexible and disordered regions in redox-sensitive PRKs.** $C_\alpha$ trace of (A), *Cr*PRK and (B), *At*PRK. The trace thickness is proportional to the atomic B factor. *At*PRK shows a higher number of flexible regions compared to *Cr*PRK. The residues belonging to the flexible regions are reported.

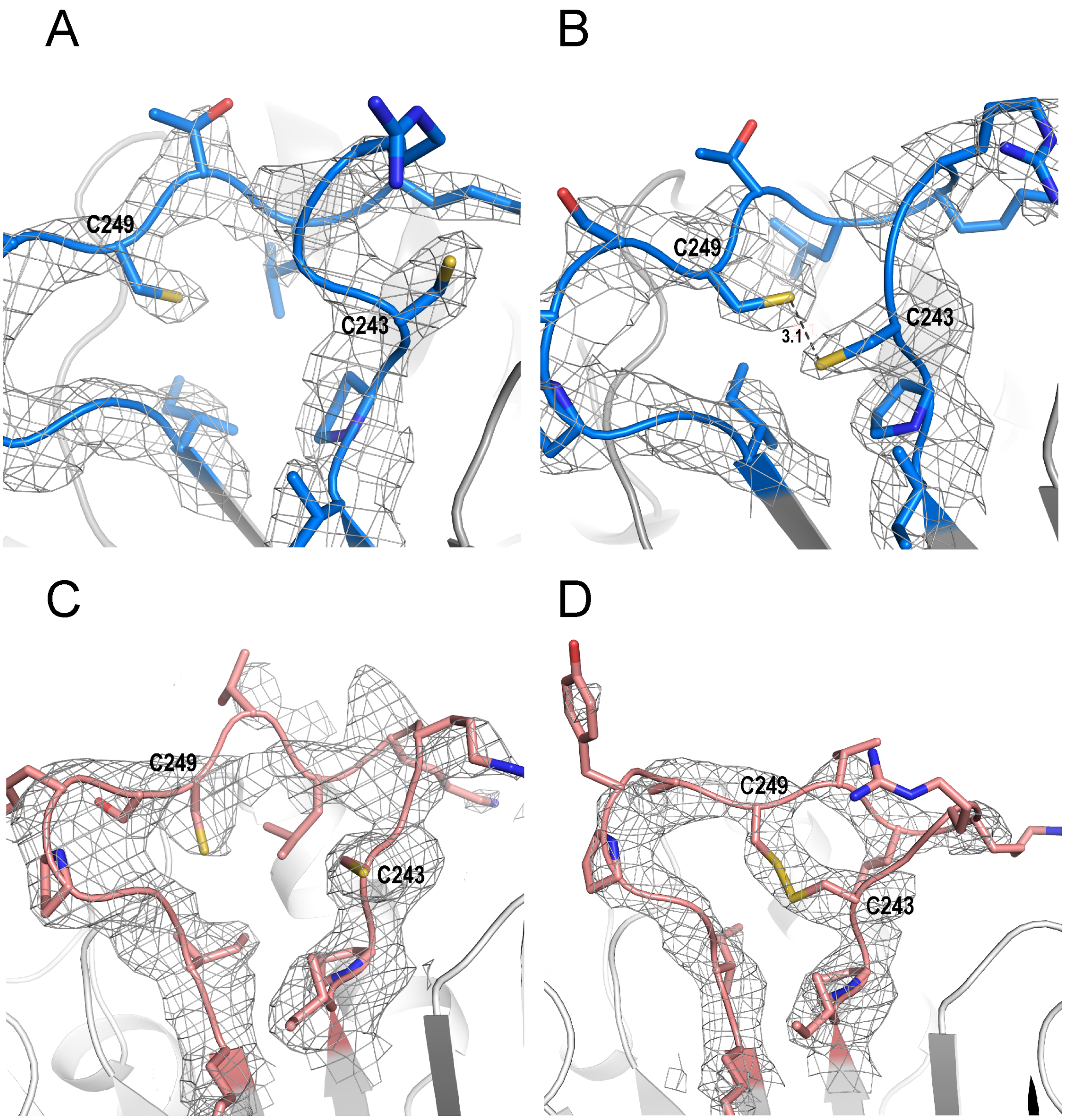


**Fig. S12. Electron density of the C-terminal cysteine pair.** $2F_o$-$F_c$ electron density map contoured at 1.2σ and associated to Cys243 and Cys249 in (A), *Cr*PRK subunit A (B), *Cr*PRK subunit B (C), *At*PRK subunit A and (D), *At*PRK subunit B. In *Cr*PRK, both C-terminal cysteines are reduced even if in subunit B the thiol groups are only 3 Å apart. A disulfide bond is instead observed in subunits B of *At*PRK, while the thiol groups of the other cysteine pair (subunit A) lie very distantly.

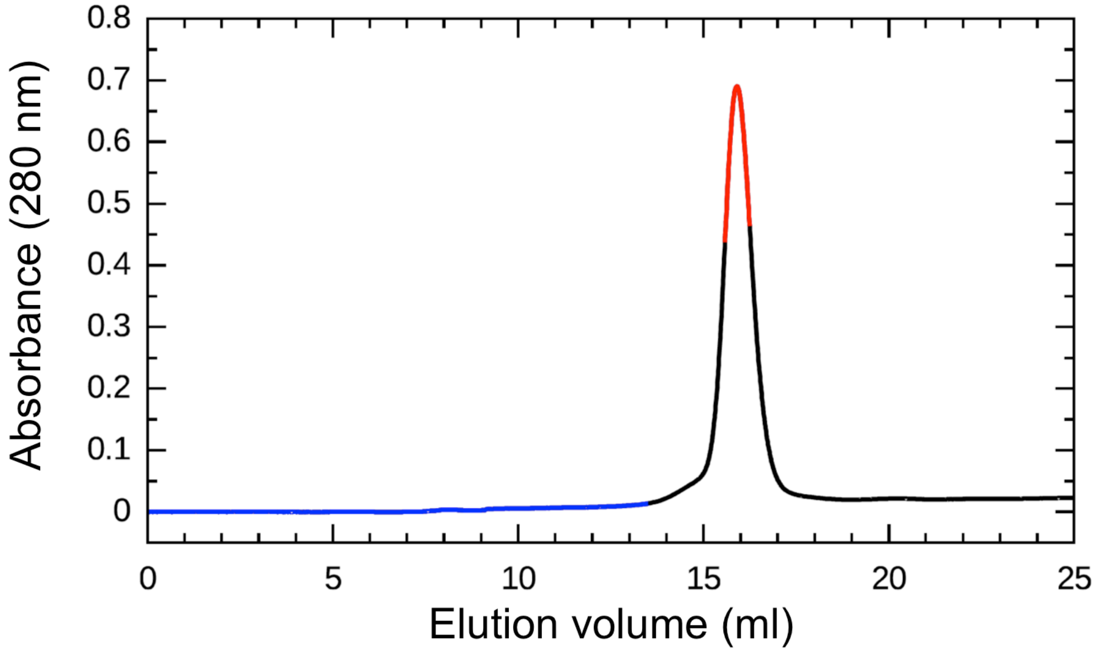


**Fig. S13. UV trace of the chromatogram profile of *Cr*PRK analyzed in SEC-SAXS mode.** Frames collected as buffer (from 0 to 14 ml) are highlighted in blue; frames collected as protein (from 14 to 17.5 ml) are highlighted in black; frames highlighted in red have been used for SAXS analysis.

**Table S1. SEC-SAXS data analysis of reduced CrPRK.**

| | |
|---|---|
| Concentration (mg $ml^{-1}$) | 6.1 (injected) |
| **Structural parameters (34)** | |
| q interval for Guinier linear fit ($nm^{-1}$) | 0.12-0.38 |
| I(0) [from Guinier approximation] | 42.0 ± 0.1 |
| $R_g$ (nm) [from Guinier approximation] | 3.43 ± 0.01 |
| q interval for Fourier inversion ($nm^{-1}$) | 0.12-3.5 |
| I(0) [from P(R)] | 42.4 ± 0.06 |
| $R_g$ (nm) [from P(R)] | 3.55 ± 0.01 |
| $D_{max}$ (nm) | 11.2±0.5 |
| Porod volume estimate ($nm^3$) | 115 ± 10 |
| DAMMIN excluded volume ($nm^3$) | 138 ± 1 |
| Dry volume calculated from sequence ($nm^3$) (v=0.735 $cm^3$ $g^{-1}$) | 95 |
| **Molecular mass (kDa)** | |
| From I(0) | 70 |
| From Vc | 70 |
| From Porod invariant | 85 |
| From Porod volume (x0.625) | 72 |
| From excluded volume (x0.5) | 69 |
| From sequence | 77.8 |

**Table S2. Accessibility values for cysteine residues and a strictly conserved arginine.**

| **Residue (*Cr*/*At*)** | **ASA (Å$^2$)*** | | | |
|---|---|---|---|---|
| | *Cr*PRK | | *At*PRK | |
| | A | B | A | B |
| Cys16/15 | 14.5 | 8.5 | 7.6 | 10.6 |
| Cys55/54 | 10.6 | 10.6 | 6.3 | 9.5 |
| Cys61 | 55.9 | 58.7 | / | / |
| Cys243 | 26.6 | 37.8 | 53.4 | 14.8 |
| Cys249 | 42.4 | 31.1 | 14.8 | 50.6 |
| Arg64/63 | 51.4 | 88.7 | 82.6 | 88.2 |

*Radius of the probe solvent molecule 1.4 Å

**Table S3. Secondary structure element content in the structurally known PRKs.**

| PRK | Quaternary structure | Helix content (%) | Sheet content (%) | Other (%) |
|---|---|---|---|---|
| *C. reinhardtii* | Dimer | 31.2 | 21.9 | 46.9 |
| *A. thaliana* | Dimer | 32.3 | 21.9 | 45.8 |
| *M. hungatei* | Dimer | 50.0 | 22.2 | 27.8 |
| *R. sphaeroides* | Octamer | 41.2 | 16.9 | 41.9 |

**Table S4. X-ray (CrPRK and AtPRK) and SEC-SAXS (CrPRK) data collection parameters.**

| | *Cr*PRK | *At*PRK | *Cr*PRK SEC-mode |
|---|---|---|---|
| Detector | Pilatus 2M | ADSC Quantum Q315r | Pilatus 1M |
| Beam geometry ($mm^2$) | 0.1 × 0.1 | 0.1 × 0.1 | 0.7 × 0.7 |
| Wavelength (Å) | 1.240 | 0.940 | 0.990 |
| Capillary diameter (mm) | / | / | 1.8 |
| Sample-to-detector distance (mm) | 239.85 | 393.43 | 2872 |
| Df (°) | 0.5 | 0.7 | / |
| q* range ($nm^{-1}$) | / | / | 0.033-4.9 |
| Exposure time (s) | 5 | 5 | 1 |
| Flow (ml/min) | / | / | 0.5 |
| Temperature (K) | 100.0 | 100.0 | 277.15 |

*$q = 4\pi \sin(\theta)/\lambda$, where $2\theta$ is the scattering angle and $\lambda$ is the X-ray wavelength.

**Table S5. X-ray data collection and refinement statistics.**

| | ***Cr*PRK** | ***At*PRK** |
|---|---|---|
| *Data collection* | | |
| Unit cell (Å) | 77.68, 83.55, 133.15, 90.00, 90.00, 90.00 | 116.30, 116.30, 106.81, 90.00, 90.00, 90.00 |
| Space group | $P2_12_12_1$ | $I4_1$ |
| Resolution range* (Å) | 44.38 – 2.60 (2.72 – 2.60) | 82.23 – 2.47 (2.58 – 2.47) |
| Unique reflections | 27230 (3280) | 24824 (3107) |
| Completeness* (%) | 99.6 (100.0) | 97.6 (99.6) |
| $R_{merge}$* | 0.076 (0.770) | 0.091 (0.435) |
| $CC_{1/2}$ | 0.997 (0.743) | 0.997 (0.947) |
| I/σ(I)* | 13.6 (1.7) | 11.2 (2.3) |
| Multiplicity* | 5.1 (5.3) | 6.9 (6.5) |
| *Refinement* | | |
| Resolution range* (Å) | 39.86 – 2.60 (2.69 - 2.60) | 46.77 – 2.47 (2.57 – 2.47) |
| Reflection used* | 27162 (2666) | 24672 (2786) |
| $R/R_{free}$ | 0.227/0.262 | 0.226/0.281 |
| rmsd from ideality (Å, °) | 0.004, 0.915 | 0.011, 1.128 |
| *N° atoms* | | |
| Non-hydrogen atoms | 5360 | 5385 |
| Protein atoms | 5319 | 5355 |
| Solvent molecules | 31 | 30 |
| Hetero atoms | 10 | / |
| *B value* ($Å^2$) | | |
| Mean | 62.4 | 78.4 |

| | | |
|---|---|---|
| Wilson | 59.0 | 52.5 |
| Protein atoms | 62.3 | 78.4 |
| Solvent molecules | 56.2 | 70.3 |
| Hetero atoms | 85.5 | / |
| *Ramachandran plot* (%)[§] | | |
| Most favoured | 91.1 | 91.4 |
| Allowed | 7.4 | 6.0 |
| Disallowed | 1.5 | 2.7 |

*Values in parentheses refer to the last resolution shell

[§]As defined by MolProbity (35)

### Table S6. Proteins used for phylogeny.

| Organism | Organism - full description | Uniprot used | other Ref. | other Ref 2 | Kingdom | Phylum | Photosynthetic | Reference and/or status of the uniprot entry |
|---|---|---|---|---|---|---|---|---|
| *A. profundus* | Archaeoglobus profundus | D2REP9 | | ADB58593 | Archae | Euryarchaeota | no | Kono et al., 2017 (5) |
| *M. concilii* | Methanosaeta concilii | F4BW53 | | AEB68483 | Archae | Euryarchaeota | no | Kono et al., 2017 (5) |
| *M. hungatei* | Methanospirillum hungatei | Q2FUB5 | | PDB: 5B3F | Archae | Euryarchaeota | no | Kono et al., 2017 (5) |
| *M. marisnigri* | Methanoculleus marisnigri | A3CWW0 | | ABN57860 | Archae | Euryarchaeota | no | Kono et al., 2017 (5) |
| *M. thermophila* | Methanosaeta thermophila | A0B947 | | ABK15221 | Archae | Euryarchaeota | no | Kono et al., 2017 (5) |
| *A. boonei* | Aciduliprofundum boonei (strain DSM 19572 / T469) | B5ICQ8 | | | Archae | Euryarchaeota | no | Kono et al., 2017 (5) / protein predicted |
| *A. veneficus* | Archaeoglobus veneficus (strain DSM 11195 / SNP6) | F2KMR8 | | | Archae | Euryarchaeota | no | Kono et al., 2017 (5) / protein predicted |
| *F. placidus* | Ferroglobus placidus (strain DSM 10642 / AEDII12DO) | D3RZ92 | | | Archae | Euryarchaeota | no | Kono et al., 2017 (5) / protein predicted |
| *M. boonei* | Methanoregula boonei (strain DSM 21154 / JCM 14090 / 6A8) | A7I4Y6 | | | Archae | Euryarchaeota | no | Kono et al., 2017 (5) / protein predicted |
| *M. harundinacea* | Methanosaeta harundinacea (strain 6Ac) | G7WL21 | | | Archae | Euryarchaeota | no | Kono et al., 2017 (5) / protein predicted |
| *M. limicola* | Methanoplanus limicola DSM 2279 | H1YX11 | | | Archae | Euryarchaeota | no | Kono et al., 2017 (5) / protein predicted |
| *M. liminatans* | Methanofollis liminatans DSM 4140 | J0SAT8 | | | Archae | Euryarchaeota | no | Kono et al., 2017 (5) / protein predicted |
| *M. palustris* | Methanosphaerula palustris (strain ATCC BAA-1556 / DSM 19958 / E1-9c) | B8GGS2 | | | Archae | Euryarchaeota | no | Kono et al., 2017 (5) / protein predicted |
| *M. petrolearia* | Methanolacinia petrolearia (strain DSM 11571 / OCM 486 / SEBR 4847) (Methanoplanus petrolearius) | E1RI35 | | | Archae | Euryarchaeota | no | Kono et al., 2017 (5) / protein predicted |
| *Methanolinea sp* | Methanolinea sp. SDB | A0A0Q0VL15 | | | Archae | Euryarchaeota | no | Kono et al., 2017 (5) / protein predicted |
| *A. ferrooxidans* | Acidithiobacillus ferrooxidans (strain ATCC 23270 / DSM 14882 / CIP 104768 / NCIMB 8455) (Ferrobacillus ferrooxidans (strain ATCC 23270)) | B7J555 | | | Bacteria | Acidithiobacillia | no | Kono et al., 2017 (5) / protein inferred from homology |
| *R. sphaeroides* | Rhodobacter sphaeroides | AAA26113 | P12033 | PDB: 1A7J | Bacteria | α-Proteobacteria | yes (purple non-sulfur) | Hallenbeck and Kaplan, 1987 (36) |
| *N. vulgaris* | Nitrobacter vulgaris | P37100 | | | Bacteria | α-Proteobacteria | no | *inferred by homology* |
| *R. meliloti* | Rhizobium meliloti (strain 1021) (Ensifer meliloti) (Sinorhizobium meliloti) | P58347 | | | Bacteria | α-Proteobacteria | no | *inferred by homology* |
| *S. medicae* | Sinorhizobium medicae (strain WSM419) (Ensifer medicae) | P56887 | | | Bacteria | α-Proteobacteria | no | *inferred by homology* |
| *R. palustris DX* | Rhodopseudomonas palustris (strain DX-1) | ADU46366 | | E6VEW3 | Bacteria | α-Proteobacteria | yes (purple non-sulfur) | Kono et al., 2017 (5) |
| *R. rubrum* | Rhodospirillum rubrum | ABC23204 | | Q2RRP1 | Bacteria | α-Proteobacteria | yes (purple non-sulfur) | Kono et al., 2017 (5) |
| *A. cryptum* | Acidiphilium cryptum (strain JF-5) | A5FWQ9 | | | Bacteria | α-Proteobacteria | no | Kono et al., 2017 (5) / protein predicted |
| *N. hamburgensis* | Nitrobacter hamburgensis (strain DSM 10229 / NCIMB 13809 / X14) | Q1QGD6 | | | Bacteria | α-Proteobacteria | no | Kono et al., 2017 (5) / protein inferred from homology |
| *R. palustris Bi* | Rhodopseudomonas palustris (strain BisA53) | Q07N64 | | | Bacteria | α-Proteobacteria | yes (purple non-sulfur) | Kono et al., 2017 (5) / protein inferred from homology |
| *X. flavus* | Xanthobacter flavus | P23015 | | | Bacteria | α-Proteobacteria | no | Meijer et al., 1991 (37) |
| *Synechocystis sp* | Synechocystis sp. (strain PCC 6803 / Kazusa) | P37101 | | | Bacteria | Cyanobacteria | yes | Experimental evidence at transcript level |
| *S. elongatus* | Synechococcus elongatus PCC 7942 | Q31PL2 | | | Bacteria | Cyanobacteria | yes | Kobayashi et al., 2003 (38) |
| *A. variabilis* | Anabaena variabilis (strain ATCC 29413 / PCC 7937) | Q3MF31 | | | Bacteria | Cyanobacteria | yes | Kono et al., 2017 (5) / protein inferred from homology |
| *G. violaceus* | Gloeobacter violaceus (strain PCC 7421) | Q7NI87 | | | Bacteria | Cyanobacteria | yes | Kono et al., 2017 (5) / protein inferred from homology |
| *T. elongatus* | Thermosynechococcus elongatus (strain BP-1) | Q8DHN2 | | | Bacteria | Cyanobacteria | yes | Kono et al., 2017 (5) / protein inferred from homology |
| *Cyanothece sp* | Cyanothece sp. (strain PCC 7424) (Synechococcus sp. (strain ATCC 29155)) | B7KI62 | | | Bacteria | Cyanobacteria | yes | Kono et al., 2017 (5) / protein predicted |
| *G. kilaueensis* | Gloeobacter kilaueensis JS1 | U5QBM9 | | | Bacteria | Cyanobacteria | yes | Kono et al., 2017 (5) / protein predicted |
| *M. aeruginosa* | Microcystis aeruginosa PCC 9701 | I4INY9 | | | Bacteria | Cyanobacteria | yes | Kono et al., 2017 (5) / protein predicted |
| *N. spumigena* | Nodularia spumigena CCY9414 | A0ZEU1 | | | Bacteria | Cyanobacteria | yes | Kono et al., 2017 (5) / protein predicted |
| *E. amylovora* | Erwinia amylovora | E5BA03 | | | Bacteria | γ-Proteobacteria | no | *inferred by homology* |
| *S. flexneri* | Shigella flexneri | P0AEX6 | | | Bacteria | γ-Proteobacteria | no | *inferred by homology* |
| *M. capsulatus* | Methylococcus capsulatus (strain ATCC 33009 / NCIMB 11132 / Bath) | Q602L2 | | | Bacteria | γ-Proteobacteria | no | Kono et al., 2017 (5) / protein predicted |
| *A. vinosum* | Allochromatium vinosum (strain ATCC 17899 / DSM 180 / NBRC 103801 / NCIMB 10441 / D) (Chromatium vinosum) | D3RP02 | | | Bacteria | γ-Proteobacteria | yes (purple sulfur) | Kono et al., 2017 (5) / protein inferred from homology |
| *P. luminescens* | Photorhabdus luminescens (Xenorhabdus luminescens) | A0A1B8YKC5 | | | Bacteria | γ-Proteobacteria | no | Kono et al., 2017 (5) / protein inferred from homology |
| *P. profundum* | Photobacterium profundum (strain SS9) | Q6LVE1 | | | Bacteria | γ-Proteobacteria | no | Kono et al., 2017 (5) / protein inferred from homology |
| *C. M. oxyfera* | Candidatus Methylomirabilis oxyfera | D5MHD4 | | | Bacteria | unclassified | no | Kono et al., 2017 (5) / protein inferred from homology |
| *T. denitrificans* | Thiobacillus denitrificans (strain ATCC 25259) | Q3SG56 | | | Bacteria | β-Proteobacteria | no | Kono et al., 2017 (5) / protein predicted |
| *C. necator* | Cupriavidus necator | P19923 | | | Bacteria | β-Proteobacteria | no | Kossmann et al., 1989 (39) |
| *E. gracilis* | Euglena gracilis | Q24LT0 | | | Excavates | Euglenozoa | yes | Experimental evidence at transcript level |
| *G. theta* | Guillardia theta | Q24LT4 | | | Hacrobia# | Cryptophyta | yes | Experimental evidence at transcript level |
| *D. lutheri* | Diacronema lutheri (Monochrysis lutheri) **Pavlova lutheri** | Q24LT5 | | | Hacrobia# | Haptophyceae | yes | Experimental evidence at transcript level |
| *P. parvum* | Prymnesium parvum | Q24LS8 | | | Hacrobia# | Haptophyceae | yes | *Predicted protein* |
| *C. variabilis* | Chlorella variabilis | E1ZF27 | | | Viridiplantae | Chlorophyte | yes | *inferred by homology* |
| *M. commoda* | Micromonas commoda | C1FFW3 | | | Viridiplantae | Chlorophyte | yes | *inferred by homology* |
| *O. tauri* | Ostreococcus tauri | A0A090N369 | | | Viridiplantae | Chlorophyte | yes | *inferred by homology* |
| *V. carteri* | Volvox carteri f. nagariensis | D8TRR7 | | | Viridiplantae | Chlorophyte | yes | *Predicted protein* |
| *C. reinhardtii* | Chlamydomonas reinhardtii | P19824 | A8IYP4 | AAF36402 | Viridiplantae | Chlorophyte | yes | Roesler and Ogren, 1990 (40) |
| *G. sulphuraria* | Galdieria sulphuraria | M2X324 | | | Viridiplantae | Rhodophyta | yes | Kono et al., 2017 (5) / protein predicted |
| *P. sativum* | Pisum sativum | P93681 | | | Viridiplantae | Streptophyta | yes | Experimental evidence at transcript level |
| *P. sitchensis* | Picea sitchensis (Sitka spruce) (Pinus sitchensis) | C0PS66 | | | Viridiplantae | Streptophyta | yes | Experimental evidence at transcript level |
| *P. patens* | Physcomitrella patens | A9TRN4 | | | Viridiplantae | Streptophyta | yes | *inferred by homology* |
| *P. trichocarpa* | Populus trichocarpa (Western balsam poplar) (Populus balsamifera subsp. trichocarpa) | B9GZT5 | | | Viridiplantae | Streptophyta | yes | *inferred by homology* |
| *S. moellendorf.* | Selaginella moellendorffii | D8SU48 | | | Viridiplantae | Streptophyta | yes | *inferred by homology* |
| *M. crystallinum* | Mesembryanthemum crystallinum | P27774 | | | Viridiplantae | Streptophyta | yes | Michalowski et al., 1992 (41) |
| *S. oleracea* | Spinacia oleracea | P09559 | | CAA30499 | Viridiplantae | Streptophyta | yes | Roesler and Ogren, 1990 (40) |
| *T. aestivum* | Triticum aestivum | P26302 | | | Viridiplantae | Streptophyta | yes | Roesler and Ogren, 1990 (40) |
| *A. thaliana* | Arabidopsis thaliana | P25697 | | | Viridiplantae | Streptophyta | yes | This Study and Roesler and Ogren, 1990 (40) |
| *L. polyedrum* | Lingulodinium polyedrum | Q24LT3 | | | SAR-Alveolates* | Dinoflagellate | yes | Experimental evidence at transcript level |
| *P. lunula* | Pyrocystis lunula | Q24LT2 | | | SAR-Alveolates* | Dinoflagellate | yes | Experimental evidence at transcript level |
| *B. natans* | Bigelowiella natans | Q24LS7 | | | SAR-Rhizaria* | Cercozoa | yes | *Predicted protein* |
| *O. sinensis* | Odontella sinensis | Q95033 | | | SAR-Stramenopiles* | Diatom | yes | Experimental evidence at transcript level |
| *P. tricornutum* | Phaeodactylum tricornutum | B5Y5F0 | | | SAR-Stramenopiles* | Diatom | yes | *inferred by homology* |
| *T. pseudonana* | Thalassiosira pseudonana | B8BZ40 | | | SAR-Stramenopiles* | Diatom | yes | *inferred by homology* |

*Member of the SAR group (42) ; #Member of the Hacrobia Kingdom (43)

phénomènes ultramicroscopiques. Ann Phys 12:161–237.

14. Porod G (1982) General theory. *Small-Angle X-Ray Scattering*, eds Glatter O, Kratky O (Academic Press, London).
15. Rambo RP, Tainer JA (2013) Accurate assessment of mass, models and resolution by small-angle scattering. Nature 496(7446):477–481.
16. Svergun DI, Petoukhov MV, Kochet MH (2001) Determination of domain structure of proteins from X-ray solution scattering. Biophys J 80(6):2946–2953.
17. Volkov VV, Svergun DI (2003) Uniqueness of ab initio shape determination in small-angle scattering. J Appl Crystallogr 36:860–864.
18. Taylor TC, Andersson I (1997) Structure of a product complex of spinach Ribulose-1,5-Bisphosphate Carboxylase/Oxygenase. Biochemistry 36(13):4041-4046.
19. Fermani S, et al. (2010) The crystal structure of photosynthetic glyceraldehyde-3-phosphate dehydrogenase (isoform A4) from Arabidopsis thaliana in complex with NAD. Acta Crystallogr Sect F Struct Biol Cryst Commun 66(Pt6):621–626.
20. Zaffagnini M, et al. (2014) High resolution crystal structure and redox properties of chloroplastic triosephosphate isomerase from Chlamydomonas reinhardtii. Mol Plant 7(1):101-120.
21. Chiadmi M, Navaza A, Miginiac-Maslow M, Jacquot JP, Cherfils J (1999) Redox signalling in the chloroplast: Structure of oxidized pea fructose-1,6-bisphosphate phosphatase. EMBO J 18(23):6809–6815.
22. Pasquini M, et al. (2017) Structural basis for the magnesium-dependent activation of transketolase from Chlamydomonas reinhardtii. Biochim Biophys Acta 1861(8):2132-2145.
23. Gütle DD, et al. (2016) Chloroplast FBPase and SBPase are thioredoxin-linked enzymes with similar architecture but different evolutionary histories. Proc Natl Acad Sci USA 24(113):6779–6784.
24. Kopp J, Kopriva S, Süss KH, Schulz GE (1999) Structure and mechanism of the amphibolic

enzyme D-ribulose-5-phosphate 3-epimerase from potato chloroplasts. J Mol Biol 287(4):761-771.

Nucleic Acids Res 45(D1):D313-D319.
